# Genome sequence of the model rice variety KitaakeX

**DOI:** 10.1101/653089

**Authors:** Rashmi Jain, Jerry Jenkins, Shengqiang Shu, Mawsheng Chern, Joel A. Martin, Dario Copetti, Phat Q. Duong, Nikki T. Pham, David A. Kudrna, Jayson Talag, Wendy S. Schackwitz, Anna M. Lipzen, David Dilworth, Diane Bauer, Jane Grimwood, Catherine R. Nelson, Feng Xing, Weibo Xie, Kerrie W. Barry, Rod A. Wing, Jeremy Schmutz, Guotian Li, Pamela C. Ronald

## Abstract

Here, we report the *de novo* genome sequencing and analysis of *Oryza sativa* ssp. *japonica* variety KitaakeX, a Kitaake plant carrying the rice XA21 immune receptor. Our KitaakeX sequence assembly contains 377.6 Mb, consisting of 33 scaffolds (476 contigs) with a contig N50 of 1.4 Mb. Complementing the assembly are detailed gene annotations of 35,594 protein coding genes. We identified 331,335 genomic variations between KitaakeX and Nipponbare (ssp. *japonica*), and 2,785,991 variations between KitaakeX and Zhenshan97 (ssp. *indica*). We also compared Kitaake resequencing reads to the KitaakeX assembly and identified 219 small variations. The high-quality genome of the model rice plant KitaakeX will accelerate rice functional genomics.

## Background

Rice (*Oryza sativa*) provides food for more than half of the world’s population [1] and also serves as a model for studies of other monocotyledonous species. Cultivated rice contains two major types of *O. sativa*, the *O. sativa indica/Xian* group and the *O. sativa japonica/Geng* group. Using genomic markers, two additional minor types have been recognized, the circum-Aus group and the circum-Basmati group [2].

The Kitaake cultivar (ssp. *japonica*), which originated at the northern limit of rice cultivation in Hokkaido, Japan [3], has emerged as a model for rice research [4] because it is extremely early flowering, easy to propagate, and short in stature [5]. Kitaake has been used to establish multiple mutant populations, including an RNAi mutant collection [6], T-DNA insertion collections [4], [7], and a whole-genome sequenced mutant population of KitaakeX, a Kitaake variety carrying the *Xa21* immune receptor gene (formerly called X.Kitaake) [8, 9]. Kitaake has been used to explore diverse aspects of rice biology, including flowering time [10], disease resistance [11], [12], [13], small RNA biology [14], and the CRISPR-Cas9 and TALEN technologies [15], [16].

The unavailability of the Kitaake genome sequence has posed an obstacle to the use of Kitaake in rice research. For example, analysis of a fast-neutron (FN) induced mutant population in KitaakeX [8], required the use of Nipponbare (ssp. *japonica)* as the reference. Additionally, CRISPR/Cas9 guide RNAs cannot be accurately designed for Kitaake without a complete sequence. To address these issues, we assembled a high-quality genome sequence of KitaakeX, compared its genome to the genomes of rice varieties Nipponbare and Zhenshan97 (ssp. *indica*), and identified genomic variations.

## Results

Kitaake has long been recognized as a rapid life-cycle variety [17], but it has yet to be systematically compared to other rice varieties. We compared the flowering time of KitaakeX with other sequenced rice varieties under long-day conditions (14 h light/10 h dark). Consistent with other studies, we found that KitaakeX flowers much earlier than other varieties (Fig. 1a, 1b), heading at 54 days after germination. Other rice varieties Nipponbare, 93-11 (ssp. *indica*), IR64 (ssp. *indica*), Zhenshan 97, Minghui 63 (ssp. *indica*), and Kasalath (aus rice cultivar) start heading at 134, 99, 107, 79, 125, and 84 days after germination, respectively (Fig. 1b).

**Fig. 1.**
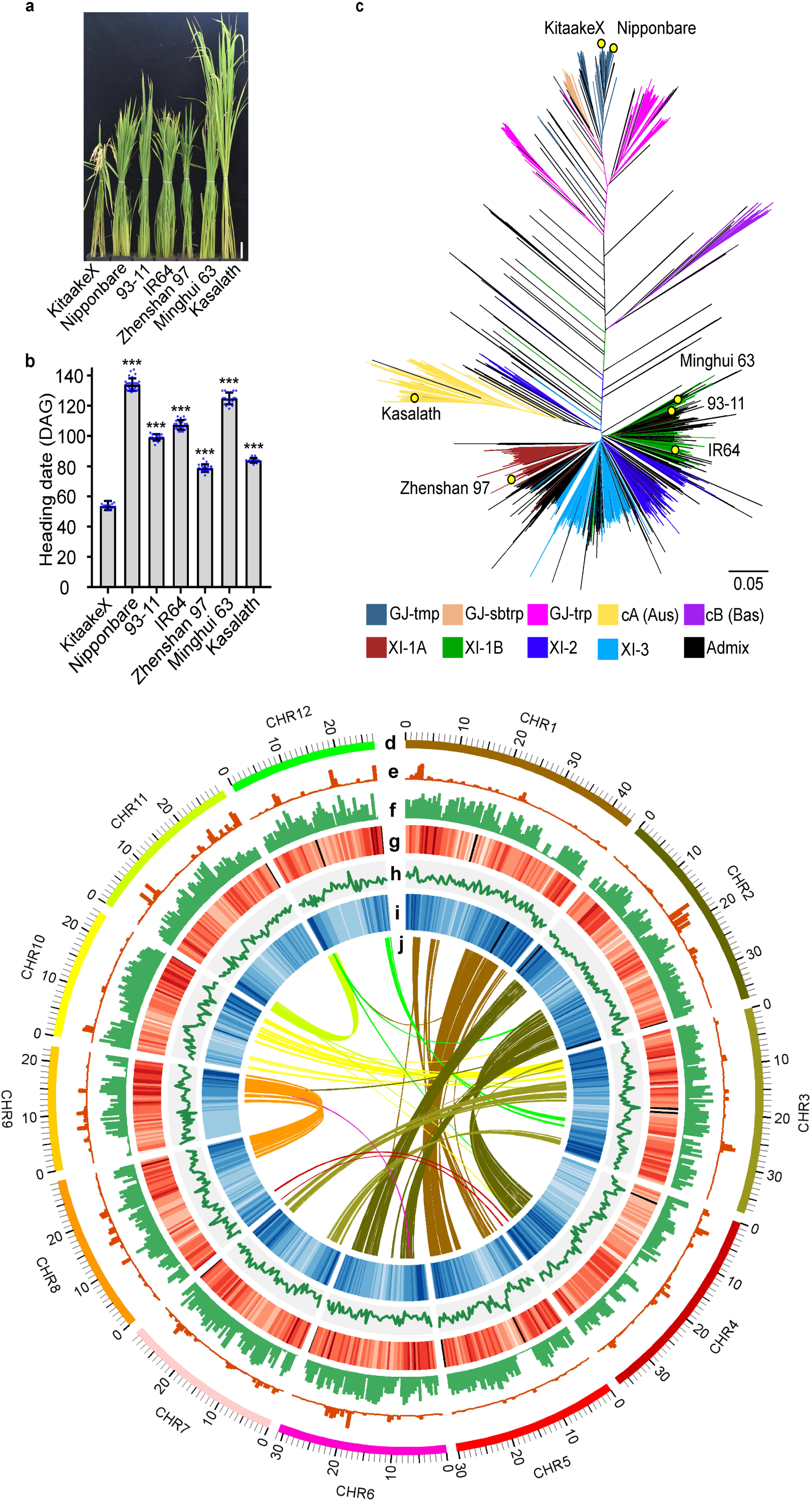
The early flowering rice variety KitaakeX. **a** KitaakeX and selected sequenced rice varieties under long-day conditions. Scale bar = 10 cm; **b** Flowering time of KitaakeX and selected rice varieties under long-day conditions. DAG, days after germination. Asterisks indicate significant differences using the unpaired Student’s *t*-test (P < 0.0001); **c** KitaakeX in the unweighted neighbor-joining tree comprising 3,010 accessions of the 3k rice genomes project and indicated varieties. It includes four XI clusters (XI-1A from East Asia, XI-1B of modern varieties of diverse origins, XI-2 from South Asia and XI-3 from Southeast Asia); three GJ clusters [primarily East Asian temperate (named GJ-tmp), Southeast Asian subtropical (named GJ-sbtrp) and Southeast Asian Tropical (named GJ-trp)]; and two groups for the mostly South Asian cA (circum-Aus) and cB (circum-Basmati) accessions, 1 group Admix (accessions that fall between major groups were classified as admixed) Branch length indicates the genetic distance between two haplotype**s; d** Circles indicate the 12 KitaakeX chromosomes represented on a Mb scale; **e**,**f** SNPs and InDels between KitaakeX and Nipponbare (**e**) and Kitaake and Zhenshan97 (**f**); **g** Repeat density; **h** GC content; **i** Gene density; **j** Homologous genes in the KitaakeX genome. Window size used in the circles is 500 kb.

We assessed how KitaakeX is related to other rice varieties using a phylogenetic approach based on the rice population structure and diversity published for 3,010 varieties [2]. The 3010 sequenced accessions were classified into nine subpopulations, most of which could be connected to geographical origins. The phylogenetic tree reveals that KitaakeX and Nipponbare are within the same subpopulation closely related (Fig 1c).

To obtain a high-quality, *de novo* genome assembly, we sequenced the KitaakeX genome using a strategy that combines short-read and long-read sequencing. Sequencing reads were collected using Illumina, 10x Genomics, PACBIO, and Sanger platforms at the Joint Genome Institute (JGI) and the HudsonAlpha Institute. The current release is version 3.0, which is a combination of a MECAT (Mapping, Error Correction and de novo Assembly Tools) PACBIO based assembly and an Illumina sequenced 10x genomics SuperNova assembly. The assembled sequence contains 377.6 Mb, consisting of 33 scaffolds (476 contigs) with a contig N50 of 1.4 Mb, covering a total of 99.67% of assembled bases in chromosomes (Table 1).

**Table 1.**
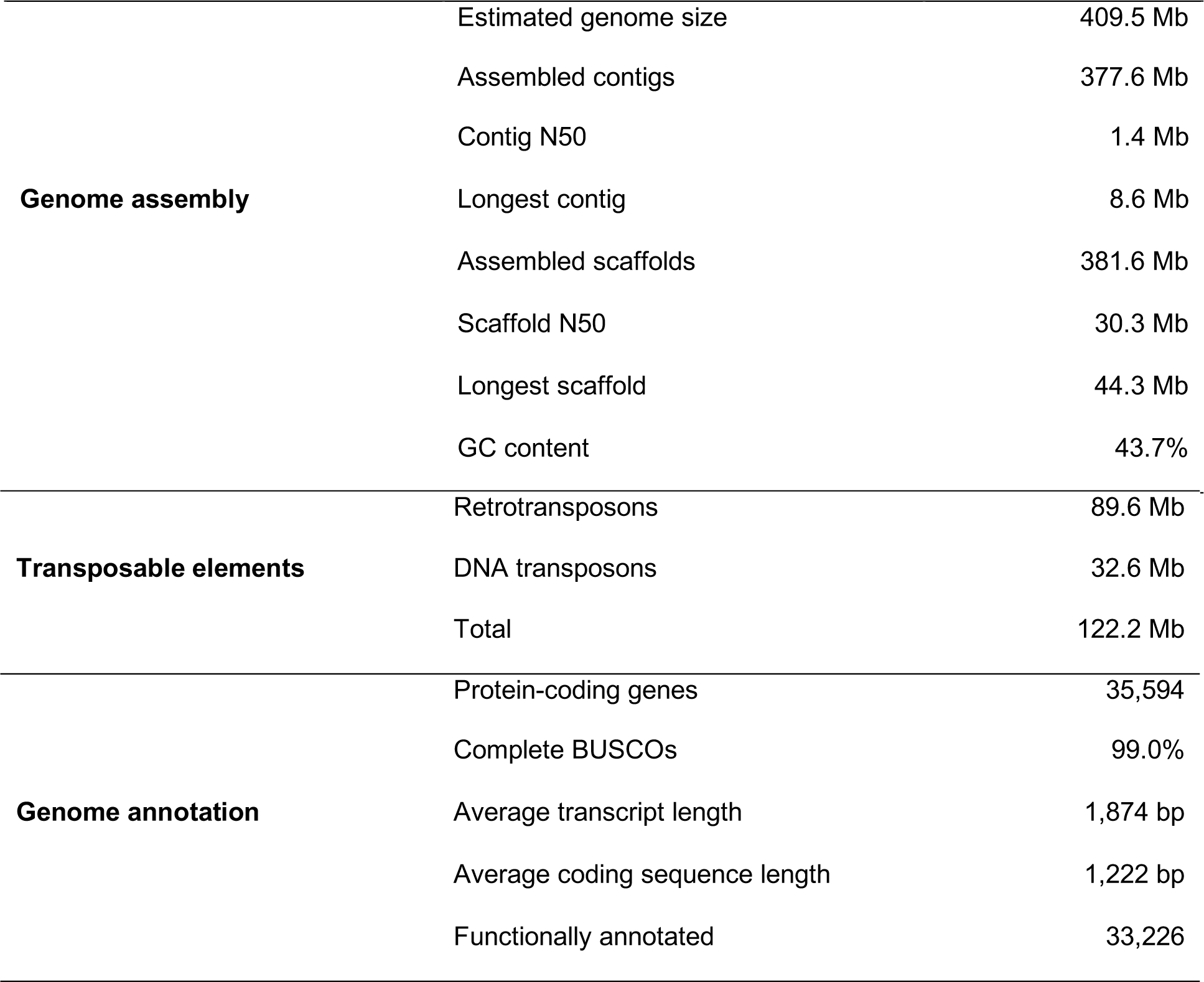
Summary of the KitaakeX genome assembly and annotation

We assessed the quality of the KitaakeX assembly for sequence completeness and accuracy. Completeness of the assembly was assessed by aligning the 34,651 annotated genes from the v7.0 Nipponbare to the KitaakeX assembly using BLAT [18]. The alignments indicate that 98.94% (34,285 of genes) genes completely aligned to the KitaakeX assembly, 0.75% (259 genes) partially aligned, and 0.31% (107 genes) were not detected. A bacterial artificial chromosome (BAC) library was constructed and a set of 346 BAC clones (9.2x clone coverage) was sequenced using PACBIO sequencing. A range of variants was detected by comparing the BAC clones to the assembly. Alignments were of high quality (<0.1% of error) in 271 clones (Additional file 1: Figure S13). Sixty BACs indicate a higher error rate (0.45% of error) due mainly to their placement in repetitive regions (Additional file 1: figure S14). Fifteen BAC clones indicate a rearrangement (10 clones) or a putative overlap on adjacent contigs (5 clones) (Additional file 1: figure S15). The overall error rate in the BAC clones is 0.09%, indicating the high quality of this assembly (for detailed information, see Additional file 1).

We predicted 35,594 protein-coding genes in the KitaakeX genome (Table 1), representing 31.5% genic space of the assembled genome size (Table 1). There is some transcriptome support for 89.5% (31,854/35,594) of the KitaakeX genes, and 81.6% (29,039/35,594) genes are fully supported by the transcriptome (Additional file 2 Table: S11). The predicted protein-coding genes are distributed unevenly across each chromosome; gene density tends to be higher toward chromosome ends (Fig. 1i). The average GC content of the genome is 43.7% (Fig. 1h, Table 1).

To assess the quality of the annotation of Kitaake genes, we compared the KitaakeX annotation to those of other completed rice genomes using the BUSCO v2 method, which is based on a set of 1,440 conserved plant genes. The results confirm 99.0% completeness of the KitaakeX genome annotation (Table1, Additional file 2: Table S7). To further evaluate the quality of the annotation, we studied the extent of conservation of functional genes in KitaakeX. We selected 291 genes (Additional file 3) from three pathways associated with stress resistance, flowering time and response to light [19], and then searched for orthologous genes in the KitaakeX genome. We found that 275 of 291 (94.5%) of the selected KitaakeX genes show greater than 90% identity with the corresponding Nipponbare genes at the protein level. Twenty-three out of the 291 show 100% identity on the genome level but not on the protein level. Of these 23 genes, the KitaakeX gene model for 16 genes has better transcriptomic evidence than the Nipponbare gene model. One of the 291 KitaakeX genes is slightly shorter than its Nipponbare ortholog due to an alternative transcript (Additional file 3). These results indicate the high quality of the annotation, and conservation between the KitaakeX and Nipponbare *japonica* rice varieties.

Using SynMap, we identified 2,469 pairs of colinear genes (88 blocks) in the KitaakeX genome (Fig. 1j). These results correlate with already published findings [20]. We used RepeatMaker and Blaster to identify transposable elements (TEs) in the KitaakeX genome, and identified 122.2 Mb of sequence corresponding to TEs (32.0% of the genome). DNA transposons account for ∼33 Mb; retrotransposons account for ∼90 Mb. The TEs belong mostly to the Gypsy and Copia retroelement families, and account for 23% of the genome (Additional file 2: Table S8), as is true in the Nipponbare and Zhenshan97 genomes [21].

We compared the genome of KitaakeX to the Nipponbare and Zhenshan97 genomes to detect genomic variations, including single nucleotide polymorphisms (SNPs), insertions and deletions under 30 bp (InDels), presence/absence variations (PAVs), and inversions using MUMmer (Kurtz, Phillippy et al. 2004). We found 331,335 variations between KitaakeX and Nipponbare (Additional file 4), and nearly 10 times as many (2,785,991) variations between KitaakeX and Zhenshan97 (Additional file 5). There are 253,295 SNPs and 75,183 InDels between KitaakeX and Nipponbare, and 2,328,319 SNPs and 442,962 InDels between KitaakeX and Zhenshan97 (Additional files 6 and Additional file 2: Table S3). With respect to SNPs in both intersubspecies (*japonica* vs. *indica*) as well as intrasubspecies (*japonica* vs. *japonica*) comparisons, transitions (Tss) (G ->A and C ->T) are about twice as abundant as transversions (Tvs) (G ->C and C ->G) (Additional file 2: Table S10). Genomic variations between KitaakeX and Nipponbare are highly concentrated in some genomic regions (Fig. 1e), but variations between KitaakeX and Zhenshan97 are spread evenly through the genome (Fig. 1f). Intersubspecies genomic variations, then, are much more extensive than intrasubspecies variations. We also detected multiple genomic inversions using comparative genomics (Additional files 4 and 5).

For variations occurring in the genic regions, we found that single-base and 3 bp (without frame shift) InDels are much more abundant than others (Additional file 7: Figure S16a), suggesting that these genetic variations have been functionally selected. We carried out detailed analysis of gene structure alterations that exist as a consequence of SNPs and InDels between KitaakeX and Nipponbare and Kitaake and Zhenshan97. Between KitaakeX and Nipponbare, we identified 2,092 frameshifts, 78 changes affecting splice-site acceptors, 71 changes affecting splice-site donors, 19 lost start codons, 161 gained stop codons, and 15 lost stop codons. In the comparison of KitaakeX to Zhenshan97, 6,809 unique genes in KitaakeX are affected by 8,640 frameshifts (Additional file 7: Figure S16b), 531 changes affecting splice-site acceptors, 530 changes affecting splice-site donors, 185 lost start codons, 902 gained stop codons and 269 lost stop codons (Additional file 7: Figure S16b).

Based on PAV analysis, we identified 456 loci that are specific to KitaakeX (Additional file 4) compared with Nipponbare. Pfam analysis of KitaakeX-specific regions revealed 275 proteins. Out of these 275 genes, 148 genes are from 19 different gene families with more than 2 genes in those regions. These gene families include protein kinases, leucine-rich repeat proteins, NB-ARC domain-containing proteins, F-box domain containing proteins, protein tyrosine kinases, Myb/SANt-like DNA binding domain proteins, transferase family proteins, xylanase inhibitor C-terminal protein, and plant proteins of unknown function (Additional file 7: Figure S16c). We identified 4589 loci specific to KitaakeX compared with Zhenshan97 (Additional file 5).

We also compared our *de novo* assembly of KitaakeX genome with Kitaake resequencing reads using an established pipeline [22]. This analysis revealed 219 small variations (200 SNPs and 19 INDELs) between the two genomes (Additional file 8). These variations affect 9 genes in KitaakeX besides the Ubi-*Xa21* transgene, including the selectable marker encoding a hygromycin B phosphotransferase on chromosome 6 (Additional file 8, Additional file 9: Figure S17).

## Discussion

In 2005 the Nipponbare genome was sequenced and annotated to a high-quality level (International Rice Genome Sequencing and Sasaki 2005). Since that time, it has served as a reference genome for many rice genomic studies [23]. Despite its use, the long life cycle of Nipponbare makes it time-consuming for most genetic analyses. Here we report the *de novo* assembly and annotation of KitaakeX, an early-flowering rice variety with a rapid life cycle that is easy to propagate under greenhouse conditions. We predict that KitaakeX contains 35,594 protein-coding genes, comparable to the published genomes (39,045 for Nipponbare and 34,610 for Zhenshan97) (Additional file 4 and Additional file 5). The availability of a high-quality genome and annotation for KitaakeX will be useful for associating traits of interest with genetic variations, and for identifying the genes controlling those traits.

We identified 219 SNPs and InDels between the KitaakeX and Kitaake genomes. These variations may have resulted from somatic mutations that arose during tissue culture and regeneration, or they may be spontaneous mutations [24]. For rice, 150 mutations are typically induced during tissue culture and 41 mutations occur spontaneously per three generations [24]. These numbers are consistent with the independent propagation of KitaakeX and Kitaake over approximately 10 generations in the greenhouse.

The KitaakeX genome will be useful for variety of studies. For example, we recently published the whole genome sequences of 1,504 FN-mutated KitaakeX rice lines [22]. Mutations were identified by aligning reads of the KitaakeX mutants to the Nipponbare reference genome [8]. On average, 97% of the Nipponbare genome is covered by the KitaakeX reads. However, in some regions, the KitaakeX genome diverges from Nipponbare to such an extent that no variants can be confidently identified. These appear either as gaps in coverage or as regions containing a concentration of natural variations between KitaakeX and Nipponbare. We can now use the KitaakeX sequence as the direct reference genome and detect mutations in highly variable regions. This approach will simplify analysis and increase confidence in the identification of FN-induced mutations.

## Conclusions

The *de novo* assembly of the KitaakeX genome serves as a useful reference genome for the model rice variety Kitaake and will facilitate investigations into the genetic basis of diverse traits critical for rice biology and genetic improvement.

## Methods

### Plant Growth conditions

Rice seeds were germinated on 1/2x MS (Murashige and Skoog) medium. Seedlings were transferred to a greenhouse and planted 3 plants/pot during the springtime (Mar. 2, 2017) in Davis, California. The light intensity was set at approximately 250 μmol m− 2 s− 1. The day/night period was set to 14/10 h, and the temperature was set between 28 and 30 °C [25]. Rice plants were grown in sandy soil supplemented with nutrient water. The day when the first panicle of the plant emerged was recorded as the heading date for that plant. Kasalath seeds were received later, and the heading date was recorded in the same way. The experiment was repeated in winter.

### Construction of a phylogenetic tree

We obtained 178,496 evenly distributed SNPs by dividing the genome into 3.8 kb bins and selecting one or two SNPs per bin randomly according to the SNP density of the bin. Genotypes of all the rice accessions, including 3,010 accessions of the 3K Rice Genomes Project and additional noted accessions, were fetched from the SNP database RiceVarMap v2.0 [26] and related genomic data [27] and used to calculate an IBS distance matrix which was then applied to construct a phylogenetic tree by the unweighted neighbor-joining method, implemented in the R package APE [28]. Branches of the phylogenetic tree were colored according to the classification of the 3,010 rice accessions [2].

### Genome Sequencing and Assembly

High molecular weight DNA from young leaves of KitaakeX was isolated and used in sequencing. See (Additional file 1) for further details.

### Annotation of Protein-Coding Genes

To obtain high-quality annotations, we performed high throughput RNA-seq analysis of libraries from diverse rice tissues (leaf, stem, panicle, and root). Approximately 683 million pairs of 2×151 paired-end RNA-seq reads were obtained and assembled using a comprehensive pipeline PERTRAN (Shu, unpublished). Gene models were predicted by combining *ab initio* gene prediction, protein-based homology searches, experimentally cloned cDNAs/expressed-sequence tags (ESTs) and assembled transcripts from the RNA-seq data. Gene functions were further annotated according to the best-matched proteins from the SwissProt and TrEMBL databases [29] using BLASTP (E value < 10^-5^) (Additional file 11). Genes without hits in these databases were annotated as “hypothetical proteins”. Gene Ontology (GO) [30] term assignments and protein domains and motifs were extracted with InterPro [31]. Pathway analysis was derived from the best-match eukaryotic protein in the Kyoto encyclopedia of genes and genomes (KEGG) database [32] using BLASTP (E value<1.0e^-10^).

### Genome Synteny

We used SynMap (CoGe, www.genomevolution.org) to identify collinearity blocks using homologous CDS pairs with parameters according to Daccord et al [33] and visualized collinearity blocks using Circos [34].

### Repeat Annotation

The fraction of transposable elements and repeated sequences in the assembly was obtained merging the output of RepeatMasker (http://www.repeatmasker.org/, v. 3.3.0) and Blaster (a component of the REPET package) [35]. The two programs were run using nucleotide libraries (PReDa and RepeatExplorer) from RiTE-db [36] and an in-house curated collection of transposable element (TE) proteins, respectively. Reconciliation of masked repeats was carried out using custom Perl scripts and formatted in gff3 files. Infernal [37] was adopted to identify non-coding RNAs (ncRNAs) using the Rfam library Rfam.cm.12.2 [38]. Results with scores lower than the family-specific gathering threshold were removed; when loci on both strands were predicted, only the hit with the highest score was kept. Transfer RNAs were also predicted using tRNAscan-SE [39] at default parameters. Repeat density was calculated from the file that contains the reconciled annotation (Additional file 10).

### Analysis of Genomic Variations

Analysis of SNPs and InDels: We used MUMmer (version 3.23) [40] to align the Nipponbare and Zhenshan97 genomes to the KitaakeX genome using parameters -maxmatch -c 90 -l 40. To filter the alignment results, we used the delta -filter -1 parameter with the one-to-one alignment block option. To identify SNPs and InDels we used show-snp option with parameter (-Clr TH). We used snpEff [41] to annotate the effects of SNPs and InDels. Distribution of SNPs and InDels along the KitaakeX genome was visualized using Circos [34].

Analysis of PAVs and inversions: We used the show-coords option of MUMmer (version 3.23) with parameters -TrHcl to identify gap regions and PAVs above >86 bp in size from the alignment blocks. We used the inverted alignment blocks with ≥98% identity from the show-coords output file to identify inversions.

To identify genomic variations between Kitaake and KitaakeX we sequenced and compared the sequences using the established pipeline [22].

### BAC library construction

Arrayed BAC libraries were constructed using established protocols [42]. Please see Additional file 1 for further details.

## Additional files

Additional file 1: Supplemental method: Tables S1-S6, Figures S1-S15

Additional file 2: Comparison of the KitaakeX genome with other rice genomes and KitaakeX annotation; Tables S7-S11

Additional file 3: Genes used in annotation quality control

Additional file 4: Comparative genomics between KitaakeX and Nipponbare

Additional file 5: Comparative genomics between KitaakeX and Zhenshan97

Additional file 6: SNPs between KitaakeX and Zhenshan97

Additional file 7: Figure S16. Genomic variations showing gene variations between KitaakeX and Nipponbare and Zhenshan97

Additional file 8. Genomic variations between KitaakeX and Kitaake

Additional file 9: Figure S17. Position of the XA21 locus in the KitaakeX genome

Additional file 10: Repeat annotation

Additional file 11: Gene functional annotation

## ACKNOWLEDGMENTS

We thank Rick A. Rios, Maria E. Hernandez, and Natasha Brown for assistance in genomic DNA isolation and submission and seed organization. We thank Dr. Thomas W. Okita at Washington State University for providing the Kitaake seeds to PCR in 1995. These seeds were provided to Dr. Okita by Dr. Hiroyuki Ito, Akita National College of Technology, Japan. We thank Dr. Jan E. Leah at Colorado State University for seeds of Zhenshan 97, Minghui 63, IR64 and 93-11, and the USDA Dale Bumpers National Rice Research Center, Stuttgart, Arkansas for seeds of Kasalath.

## Availability of data and material

The genome sequencing reads and assembly have been deposited under GenBank under accession number PRJNA234782 and PRJNA448171 respectively. The assembly and annotation of the Kitaake genome are available at Phytozome (https://phytozome.jgi.doe.gov/pz/portal.html). The RNA-Seq reads of KitaakeX leaf, panicle, stem and root have been deposited under GenBank accession numbers SRP182736, SRP182738, SRP182741, and SRP182737 respectively. Genome sequencing reads for Kitaake have been deposited under GenBank under accession number SRP193308.

## AUTHOR CONTRIBUTIONS

RJ, G.L, M.C.R. and P.C.R conceived and initiated the study. R.J. and G.L carried out sequencing, assembly and annotation in collaboration with J.J., S.S., D.A.K., J.T., D.D., D.B., J.G., D.C., K.W.B., R.A.W., N.T.P, and J.S. R.J. and G.L conducted comparative genomics. F.X and W.X contributed to phylogenetic tree. R.J., P.C.R, M.S C, G.L and C.R.N. wrote the manuscript. All authors read and approved the final manuscript.

## Ethics approval and consent to participate

Not applicable

## Consent to Publication

Not applicable

