## Additional file 1 for "Genome sequence of the model rice variety KitaakeX"

**Supplemental Methods:**

***Sequencing*:** We sequenced *Oryza sativa* (var. Kitaake) using a whole genome shotgun sequencing strategy and standard sequencing protocols. Sequencing reads were collected using Illumina, PACBIO, and Sanger platforms. Illumina, PACBIO, and Sanger reads were sequenced at the Department of Energy (DOE) Joint Genome Institute (JGI) in Walnut Creek, California and the HudsonAlpha Institute in Huntsville, Alabama. Sanger BACs were sequenced using an ABI 3730XL capillary sequencer. Illumina reads were sequenced using the Illumina MISeq/HISeq platform, and the PACBIO reads were sequenced using the RSII platform. One BAC library was sequenced using Sanger sequencing (9.2x clone coverage). One 400bp insert 2x250 Illumina fragment library (34.1x), two 800bp insert 2x250 Illumina fragment libraries (69.3x), two 4kb insert 2x150 Illumina mate pair libraries (22.3x), and one 7kb insert 2x150 Illumina mate pair library (10.9x) were obtained for a total of 136.6x coverage. Additionally, four 10X genomics chromium 2x150 libraries (335x) were sequenced (See Table S1). Prior to assembly, all Illumina reads were screened for mitochondria, chloroplast, and Phix contamination. Reads composed of >95% simple sequence were removed. Illumina reads <75bp after trimming for adapter and quality (q<20) were removed. An additional deduplication step was performed on the Illumina mate pairs that identifies and retains only one copy of each PCR duplicate. The final read set consists of 248,531,203 reads for a total of 206.52x of high quality Illumina bases, along with 845,500,242 reads for a total of 335x high quality 10X chromium bases. For the PACBIO sequencing, 42 chips (P6C4 chemistry) were sequenced with a total yield of 32.4 Gb (81.0x), a single pass yield of 24.3 Gb (60.72x) with 22.12 Gb >5kb (See Table S2). After error correction, a total of 20.4 Gb (51.0x) was used in the assembly.

***Genome assembly and construction of pseudomolecule chromosomes:***

The current release is version 3.0, which is a combination of a MECAT assembly and a SuperNova assembly. The assembly process and methods used to combine the two assemblies to produce the version 3.0 release are detailed in this section. A subset of 109.36 million 10X genomics reads (31.5x assembled sequence coverage) was assembled using SuperNova [1] with default parameters. This produced 12,274 scaffolds (20,391 contigs), with a scaffold N50 of 1.8 Mb, 256 scaffolds larger than 100 Kb, and a total genome size of 340.6 Mb (Table S3).

An improved assembly was generated by assembling the 2,873,271 PACBIO reads (60.7x sequence coverage) using the MECAT assembler [2]. This produced 519 scaffolds (519 contigs), with a scaffold N50 of 2.0 Mb, 341 scaffolds larger than 100 Kb, and a total genome size of 381.3 Mb (Table S4). The resulting assembly was polished using Quiver [3] and scaffolded with SSPACE [4] using the existing 4KB and 7KB Illumina mate pairs. A set of 33,187 unique, non-repetitive, non-overlapping 1.0 Kb syntenic sequences was identified in the *O. sativa* (var. *japonica*) assembly, aligned to the MECAT assembly and used in the identification of misjoins. The HMG BAC ends were then used to pinpoint syntenic breakpoints. A total of 68 misjoins were identified and broken. Scaffolds were then oriented, ordered, and joined together into 12 chromosomes. A total of 394 joins were made during this process, and the chromosome joins were padded with 10,000 Ns.

A comparison of the SuperNova assembly to the MECAT assembly revealed that there were sequences contained in the SuperNova assembly that were not contained in the MECAT assembly, presumably due to the differences in the sequencing platform. The SuperNova assembly was then masked using 24mers taken from the polished MECAT assembly and the unmasked regions of the SuperNova assembly over 1.0KB were extracted and included in the final release. Six SuperNova contigs (14.5 Kb) were included as part of the V3.0 release. The combined assembly was then screened for contamination. Homozygous SNPs and InDels were corrected in the release sequence using ~60x of Illumina reads (2x250, 800bp insert) by aligning the reads using bwa mem (unpublished) and identifying homozygous SNPs and InDels with the GATK’s UnifiedGenotyper tool [5]. 99 homozygous SNPs and 15,339 homozygous InDels were corrected in the release. The final version 3.0 improved release contains 377.6 Mb of sequence, consisting of 33 scaffolds (476 contigs) with a contig N50 of 1.4 Mb and a total of 99.67% of assembled bases in chromosomes. Completeness of the euchromatic portion of the version 2.0 assembly was assessed using 34,651 annotated genes from the version 7.0 *O. sativa* (var. *japonica*) release. The genes were aligned to the assembly using BLAT [6] and alignments >=90% base pair identity and >=85% gene coverage were retained. The screened alignments indicate that 34,285 (98.94%) of the *japonica* genes aligned to the Kitaake assembly, with 259 genes indicating a partial alignment (0.75%), and 107 sequences (0.31%) indicating no alignment.

SI material:

**S1.1. Assembly Construction.** A subset of 109.36 10X genomics reads (31.5x assembled sequence coverage) was assembled using SuperNova [1] with default parameters. The additional 2,873,271 PACBIO reads were assembled using MECAT [2], and a combination of the SuperNova and MECAT assemblies was used to form the version 3.0 release.

| **Library** | **Sequencing Platform** | **Average Read/Insert Size** | **Read Number** | **Assembled Sequence Coverage (x)** |
| --- | --- | --- | --- | --- |
| HMG | Sanger | 135,000 ± 30,000 | 76,543 | 9.2 clone |
| AHZPU | Illumina | 400 ± 50 | 47,644,256 | 34.1 |
| SNNP | Illumina | 735 ± 100 | 47,039,120 | 33.9 |
| TGWO | Illumina | 717 ± 100 | 49,242,748 | 35.4 |
| ACWWN | Illumina | 4,432 ± 593 | 34,513,336 | 11.0 |
| ANUPU | Illumina | 4,179 ± 551 | 33,822,898 | 11.3 |
| AHZWG | Illumina | 7,034 ± 578 | 33,319,031 | 10.9 |
| IHQR | 10X Genomics | 400 ± 50 | 845,500,242 | 335 |
|  | PACBIO | 8,072^*^ | 2,873,271 | 60.72 |
| **Total** |  | N/A | 248,531,203 | 206.52 |

**Table S1.** Genomic libraries included in the *Oryza sativa* (var. Kitaake) genome assembly and their respective assembled sequence coverage levels in the final release.

^*^Average read length of PACBIO reads.

| **Cutoff** | **Number of Reads** | **Basepairs** | **Average Read Length** | **Coverage** |
| --- | --- | --- | --- | --- |
| 0 | 2,873,271 | 24,286,612,782 | 8,072 | 60.72x |
| 1,000 | 2,769,767 | 24,227,814,611 | 8,289 | 60.57x |
| 2,000 | 2,630,301 | 24,016,773,360 | 8,584 | 60.04x |
| 3,000 | 2,467,877 | 23,609,351,251 | 8,932 | 59.02x |
| 4,000 | 2,287,691 | 22,977,809,096 | 9,325 | 57.44x |
| 5,000 | 2,098,922 | 22,128,129,658 | 9,744 | 55.32x |
| 6,000 | 1,907,045 | 21,072,464,943 | 10,184 | 52.68x |
| 7,000 | 1,694,550 | 19,688,149,643 | 10,697 | 49.22x |
| 8,000 | 1,454,022 | 17,883,760,799 | 11,324 | 44.71x |
| 9,000 | 1,218,425 | 15,882,127,906 | 12,015 | 39.71x |
| 10,000 | 993,394 | 13,745,612,155 | 12,782 | 34.36x |
| 11,000 | 787,733 | 11,588,418,240 | 13,631 | 28.97x |
| 12,000 | 611,669 | 9,566,478,012 | 14,551 | 23.92x |
| 13,000 | 468,194 | 7,775,706,908 | 15,514 | 19.44x |
| 14,000 | 356,123 | 6,265,216,945 | 16,500 | 15.66x |
| 15,000 | 270,175 | 5,020,965,093 | 17,497 | 12.55x |
| 16,000 | 204,620 | 4,006,458,829 | 18,502 | 10.02x |
| 17,000 | 154,971 | 3,188,408,846 | 19,509 | 7.97x |
| 18,000 | 117,598 | 2,535,242,511 | 20,500 | 6.34x |
| 19,000 | 89,093 | 2,008,566,675 | 21,503 | 5.02x |

**Table S2.** PACBIO library statistics for single pass yield of the 42 chips included in the *Oryza sativa* (var. Kitaake) genome assembly and their respective assembled sequence coverage levels.

| **Minimum**  **Scaffold**  **Length** | **Number of**  **Scaffolds** | **Number of**  **Contigs** | **Scaffold Size** | **Basepairs** | **% Non-gap Basepairs** |
| --- | --- | --- | --- | --- | --- |
| 5 Mb | 9 | 1,367 | 62,333,111 | 62,197,311 | 99.78% |
| 2.5 Mb | 26 | 2,671 | 119,371,803 | 119,107,303 | 99.78% |
| 1 Mb | 93 | 5,501 | 230,425,623 | 229,884,823 | 99.77% |
| 500 Kb | 149 | 6,540 | 270,151,260 | 269,512,160 | 99.76% |
| 250 Kb | 192 | 6,990 | 285,238,462 | 284,558,662 | 99.76% |
| 100 Kb | 256 | 7,319 | 296,096,685 | 295,390,385 | 99.76% |
| 50 Kb | 308 | 7,466 | 299,917,570 | 299,201,770 | 99.76% |
| 25 Kb | 388 | 7,608 | 302,629,359 | 301,907,359 | 99.76% |
| 10 Kb | 809 | 8,099 | 308,497,622 | 307,768,622 | 99.76% |
| 5 Kb | 2,397 | 9,816 | 319,366,213 | 318,624,313 | 99.77% |
| 2.5 Kb | 5,430 | 13,095 | 329,926,754 | 329,160,254 | 99.77% |
| 1 Kb | 12,274 | 20,391 | 340,592,618 | 339,780,918 | 99.76% |
| 0 bp | 12,274 | 20,391 | 340,592,618 | 339,780,918 | 99.76% |

**Table S3**. Summary statistics of the output of the SuperNova whole genome shotgun assembly prior to integration with the PACBIO assembly. The table shows total contigs and total assembled basepairs for each set of scaffolds greater than the size listed in the left hand column.

| **Minimum**  **Scaffold**  **Length** | **Number of**  **Scaffolds** | **Number of**  **Contigs** | **Scaffold Size** | **Basepairs** | **% Non-gap Basepairs** |
| --- | --- | --- | --- | --- | --- |
| 5 Mb | 8 | 8 | 57,771,145 | 57,771,145 | 100.00% |
| 2.5 Mb | 40 | 40 | 167,464,984 | 167,464,984 | 100.00% |
| 1 Mb | 112 | 112 | 276,812,732 | 276,812,732 | 100.00% |
| 500 Kb | 192 | 192 | 335,638,702 | 335,638,702 | 100.00% |
| 250 Kb | 264 | 264 | 361,964,929 | 361,964,929 | 100.00% |
| 100 Kb | 341 | 341 | 374,712,160 | 374,712,160 | 100.00% |
| 50 Kb | 385 | 385 | 377,749,994 | 377,749,994 | 100.00% |
| 25 Kb | 453 | 453 | 380,091,105 | 380,091,105 | 100.00% |
| 10 Kb | 517 | 517 | 381,281,419 | 381,281,419 | 100.00% |
| 5 Kb | 519 | 519 | 381,299,825 | 381,299,825 | 100.00% |
| 2.5 Kb | 519 | 519 | 381,299,825 | 381,299,825 | 100.00% |
| 1 Kb | 519 | 519 | 381,299,825 | 381,299,825 | 100.00% |
| 0 bp | 519 | 519 | 381,299,825 | 381,299,825 | 100.00% |

**Table S4**. Summary statistics of the raw output of the MECAT whole genome shotgun assembly. The table shows total contigs and total assembled basepairs for each set of scaffolds greater than the size listed in the left hand column.

**S1.2 Pseudomolecule Chromosome Construction**

The MECAT assembly was polished using Quiver [3] and scaffolded with SSPACE [4] using the existing 4KB and 7KB Illumina mate pairs. A set of 33,187 non-overlapping, non-repetitive, 1.0 Kb syntenic Japonica markers was used to integrate the scaffolded MECAT assembly scaffolds into 12 chromosomes. A total of 68 misjoins were identified and broken. Scaffolds were then oriented, ordered, and joined together into 12 chromosomes. The available BAC ends (9.2x clone coverage) were used to leverage in 12 additional scaffolds that did not contain a synteny marker. A total of 394 joins were made during this process, and the chromosome joins were padded with 10,000 Ns. A total of 6 SuperNova contigs (14.5 Kb) were included in the final assembly. The final version 3.0 improved release contains 33 scaffolds (476 contigs) that cover 377.6 Mb of sequence, with a contig N50 of 1.4 Mb. A total of 99.67% of assembled bases are anchored in the 12 chromosomes. Significant telomeric sequence was identified using the (TTTAGGG)_n_ repeat, and care was taken to make sure that it was properly oriented in the production assembly. Plots of the marker placements for the 12 chromosomes are shown in Figures S1-S12.

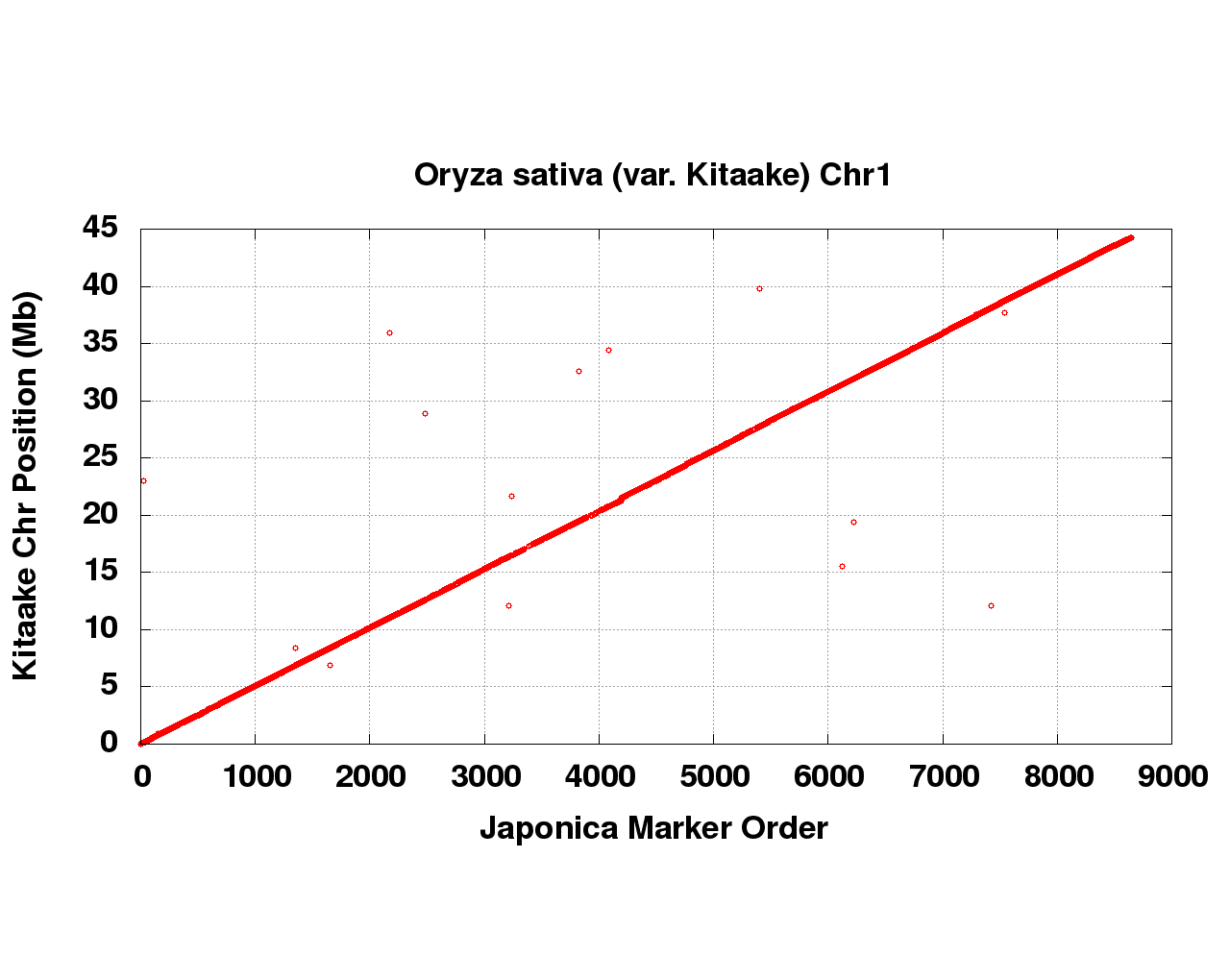

**Figure S1:** Syntenic Japonica sequence placements on the *Oryza sativa* (var. Kitaake) chromosome 1.

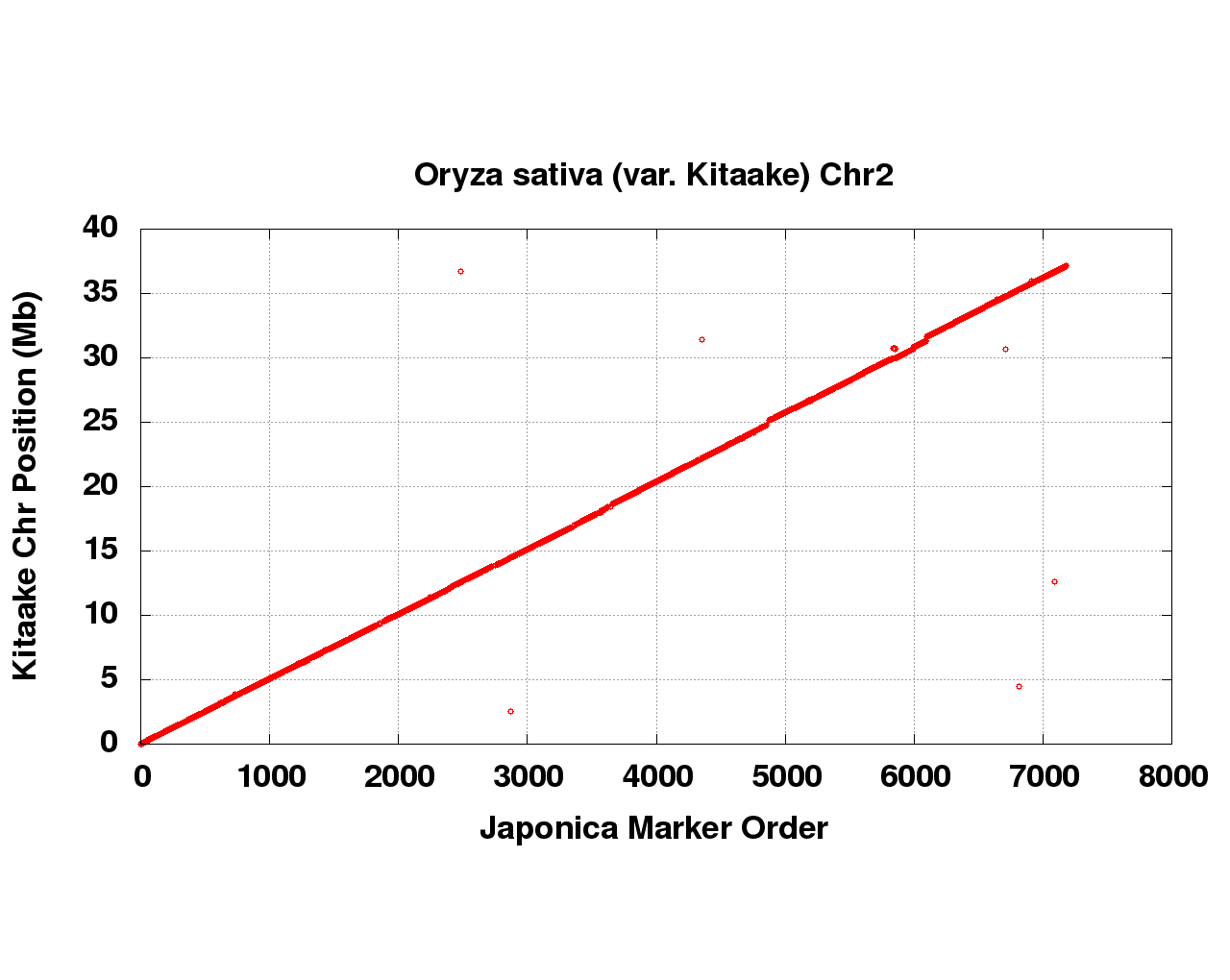

**Figure S2:** Syntenic Japonica sequence placements on the *Oryza sativa* (var. Kitaake) chromosome 2.

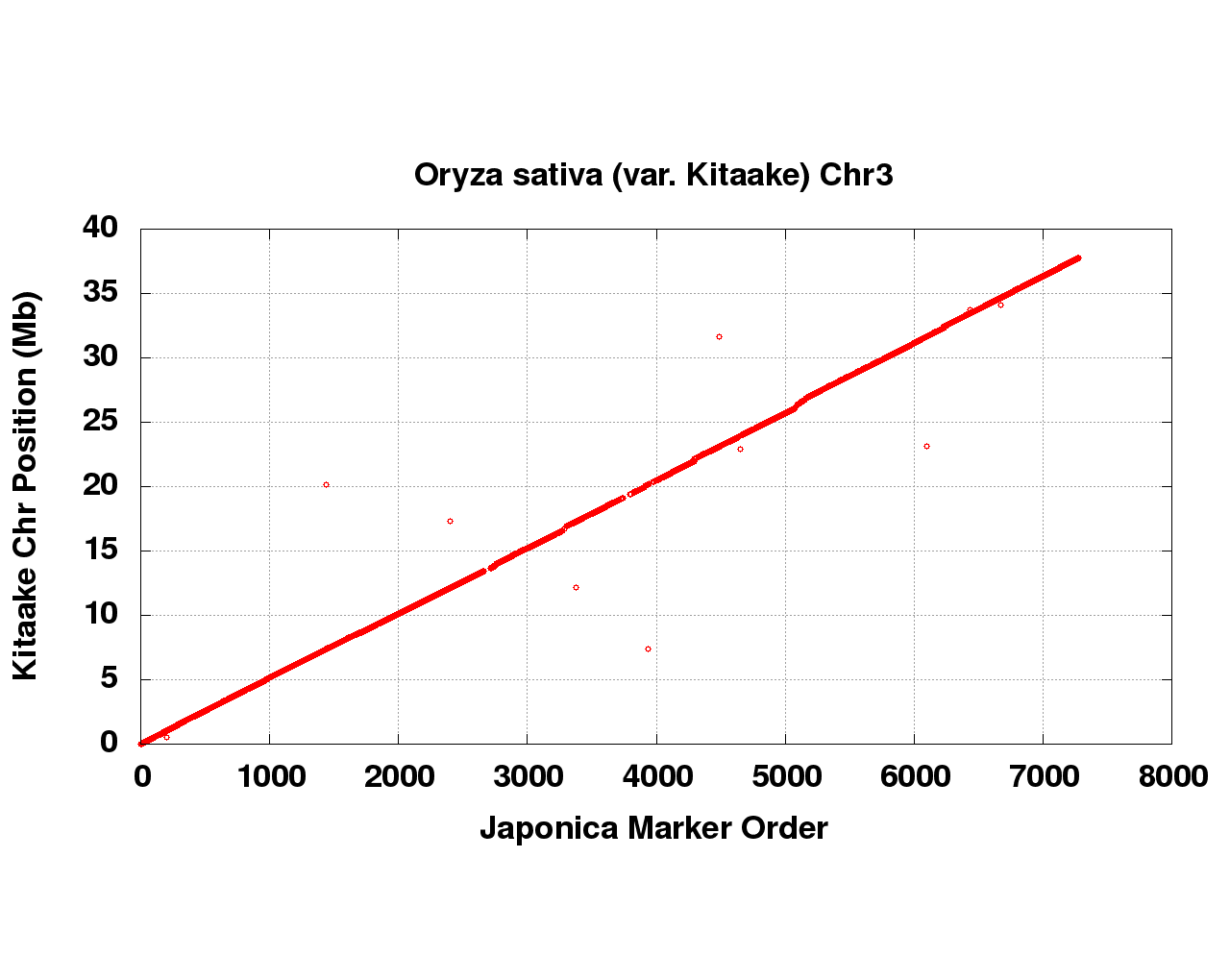

**Figure S3:** Syntenic Japonica sequence placements on the *Oryza sativa* (var. Kitaake) chromosome 3.

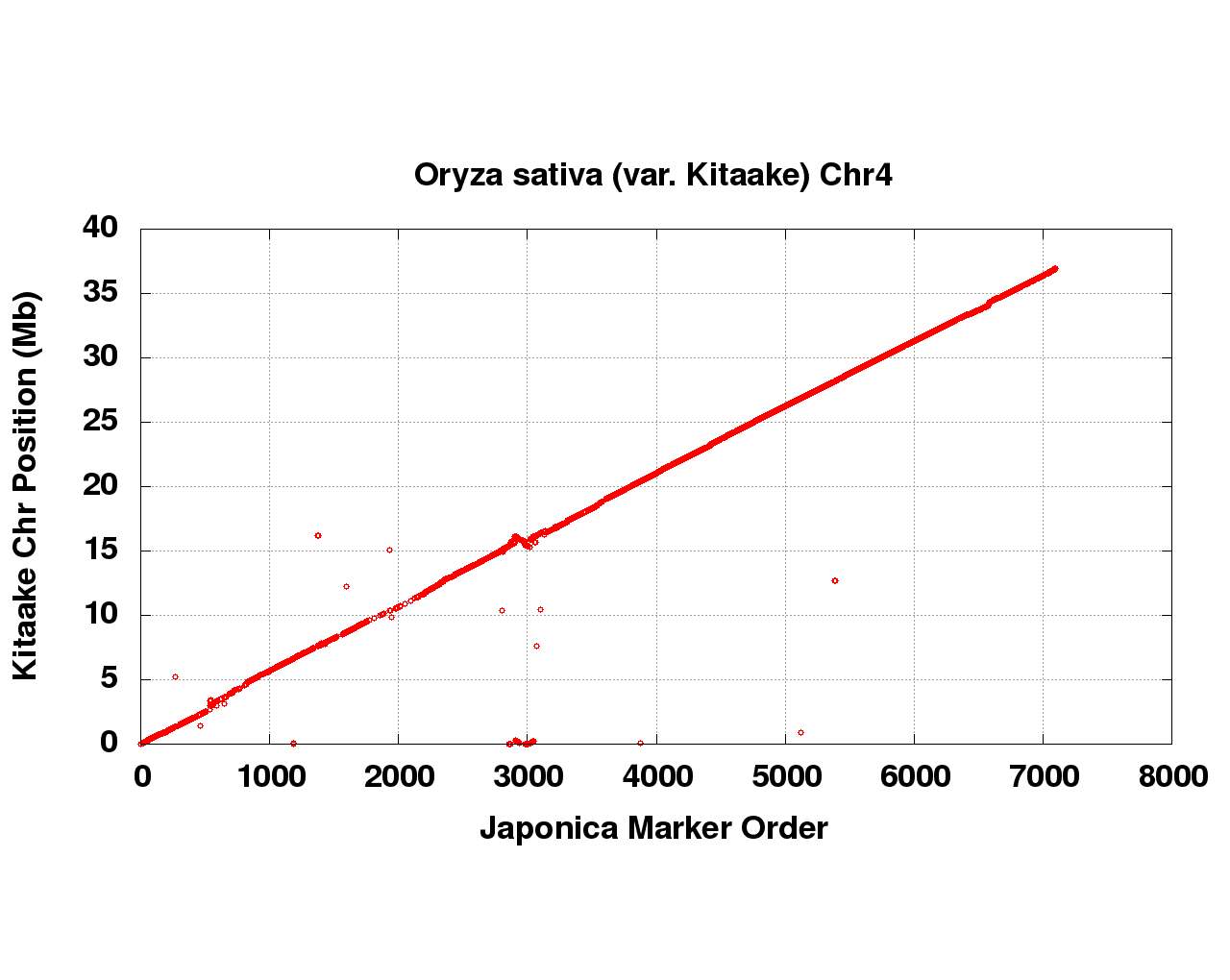

**Figure S4:** Syntenic Japonica sequence placements on the *Oryza sativa* (var. Kitaake) chromosome 4.

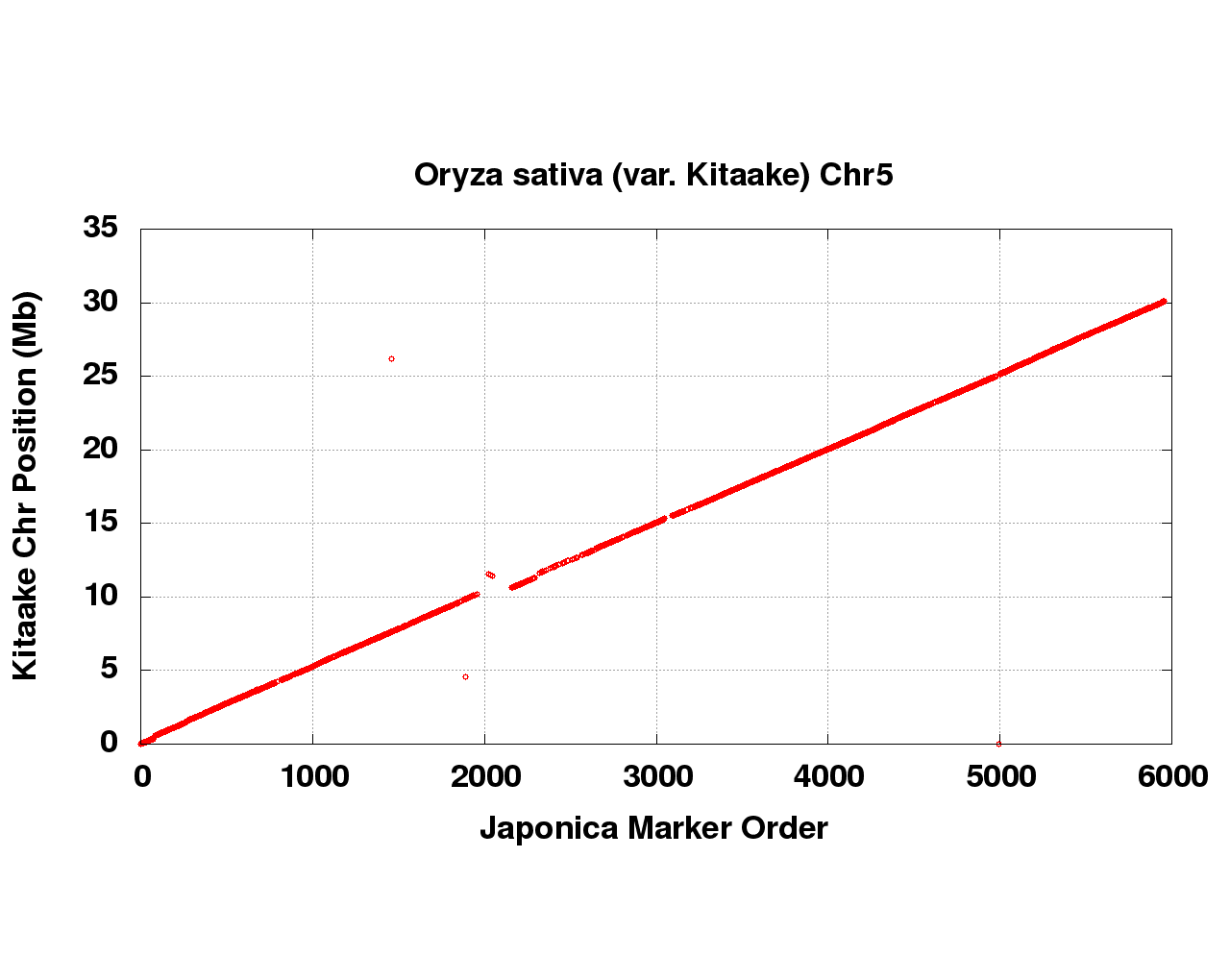

**Figure S5:** Syntenic Japonica sequence placements on the *Oryza sativa* (var. Kitaake) chromosome 5.

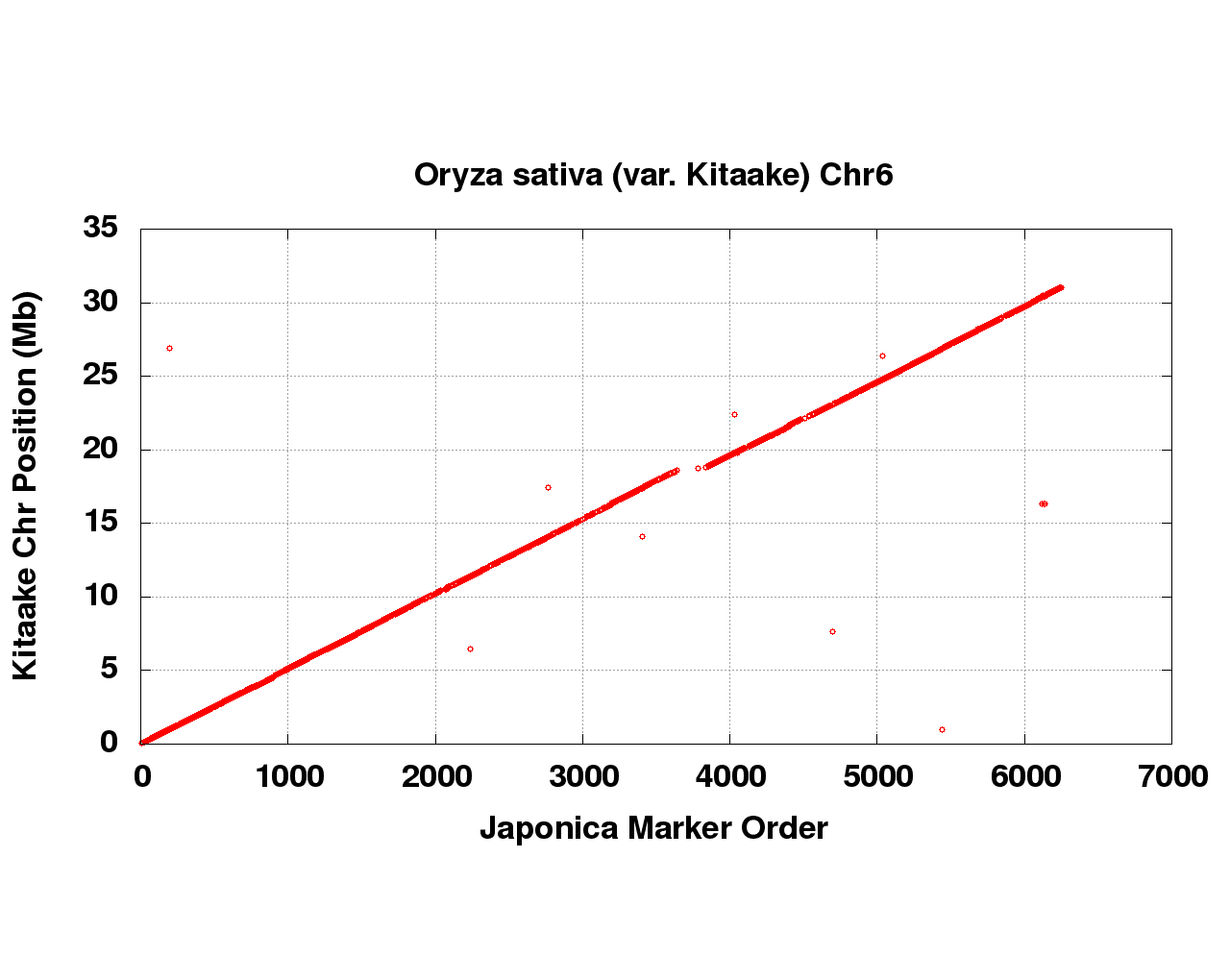

**Figure S6:** Syntenic Japonica sequence placements on the *Oryza sativa* (var. Kitaake) chromosome 6.

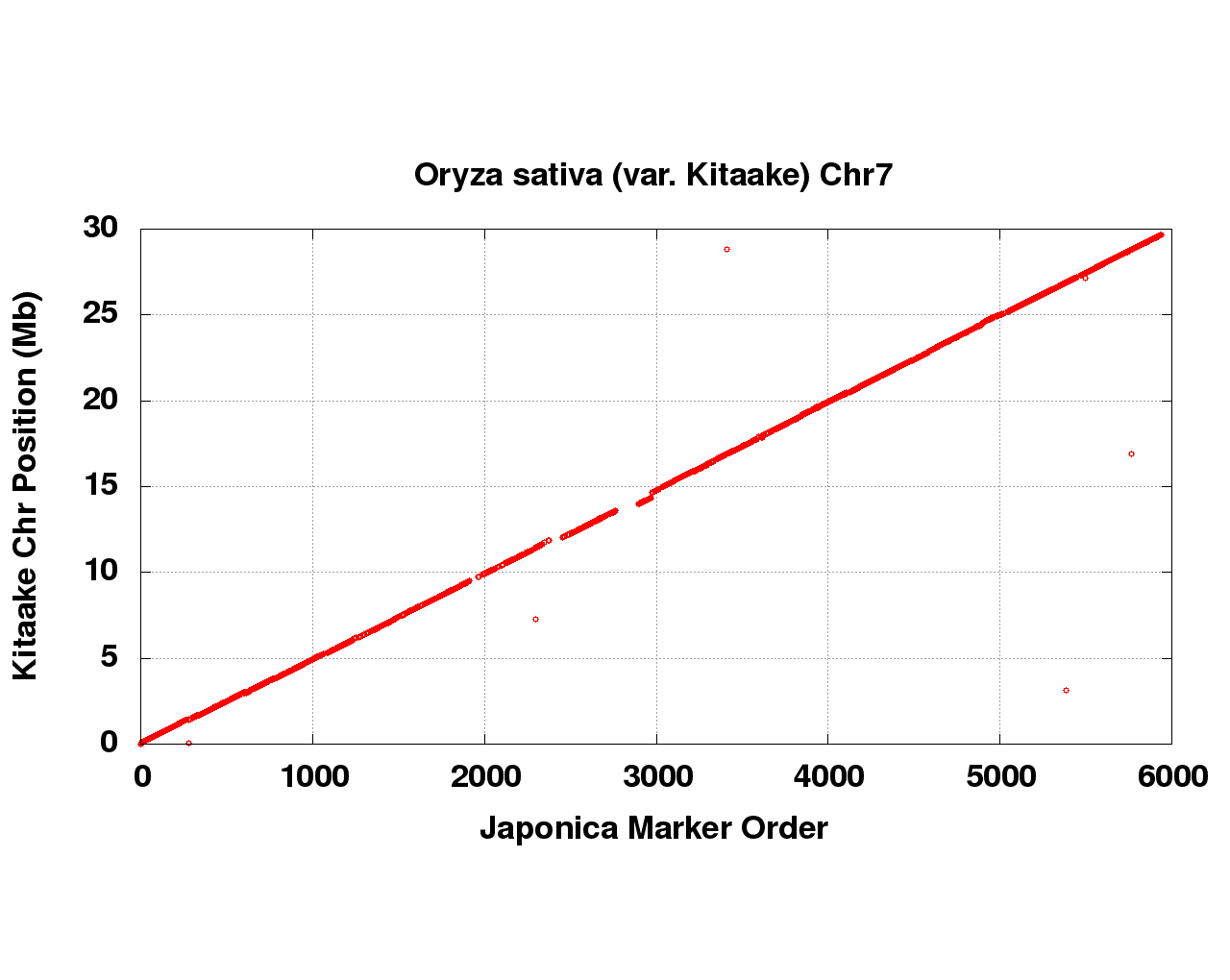

**Figure S7:** Syntenic Japonica sequence placements on the *Oryza sativa* (var. Kitaake) chromosome 7.

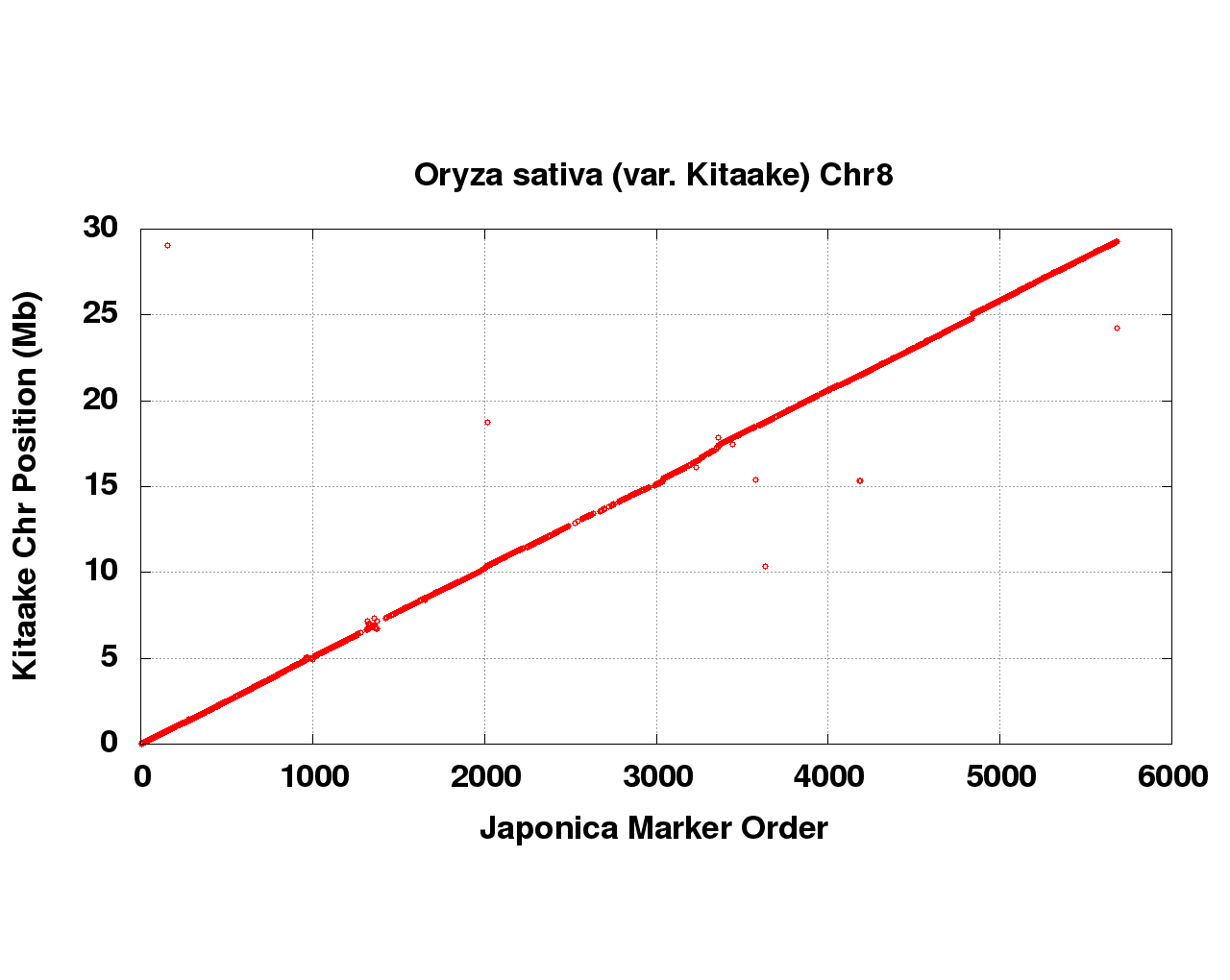

**Figure S8:** Syntenic Japonica sequence placements on the *Oryza sativa* (var. Kitaake) chromosome 8.

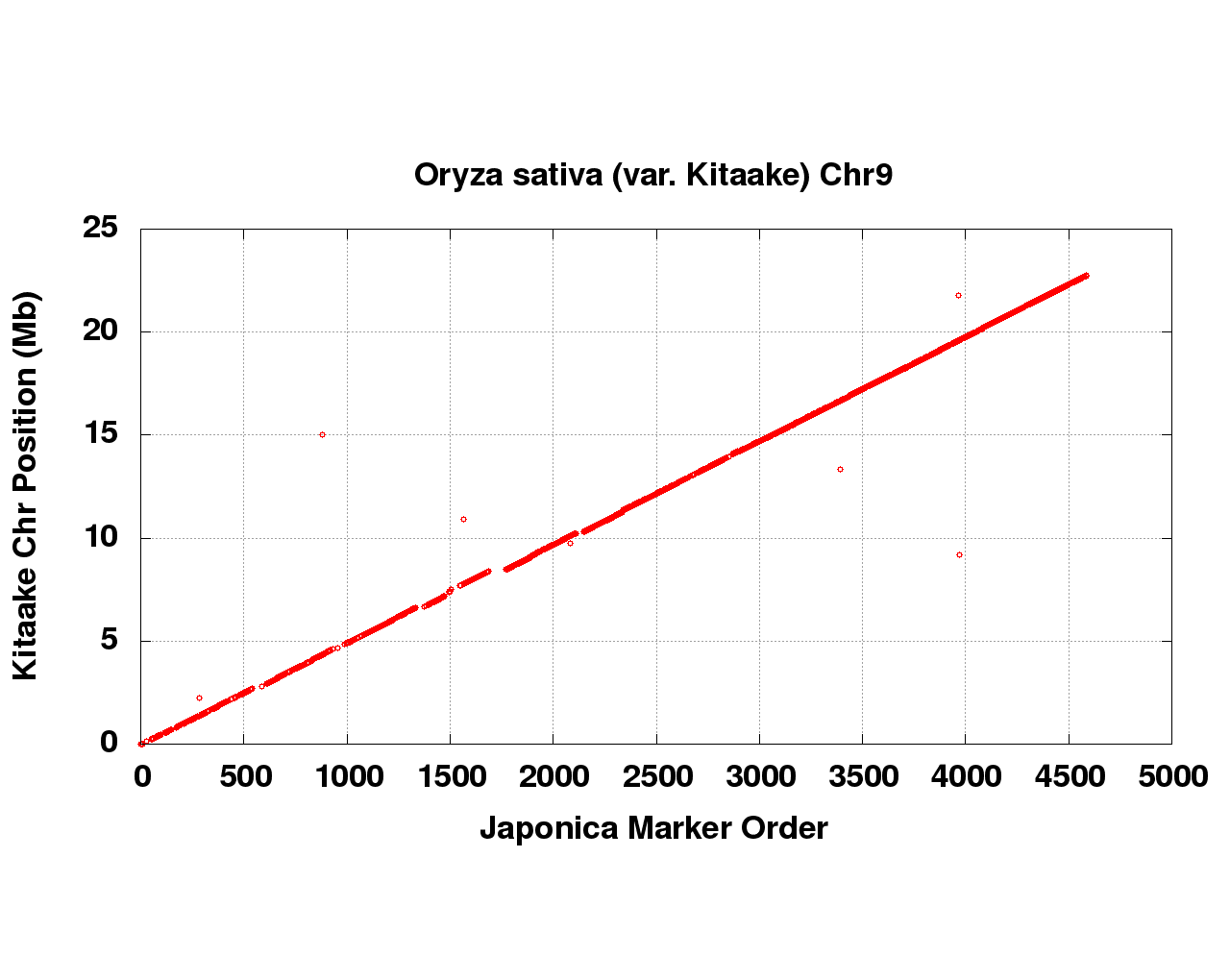

**Figure S9:** Syntenic Japonica sequence placements on the *Oryza sativa* (var. Kitaake) chromosome 9.

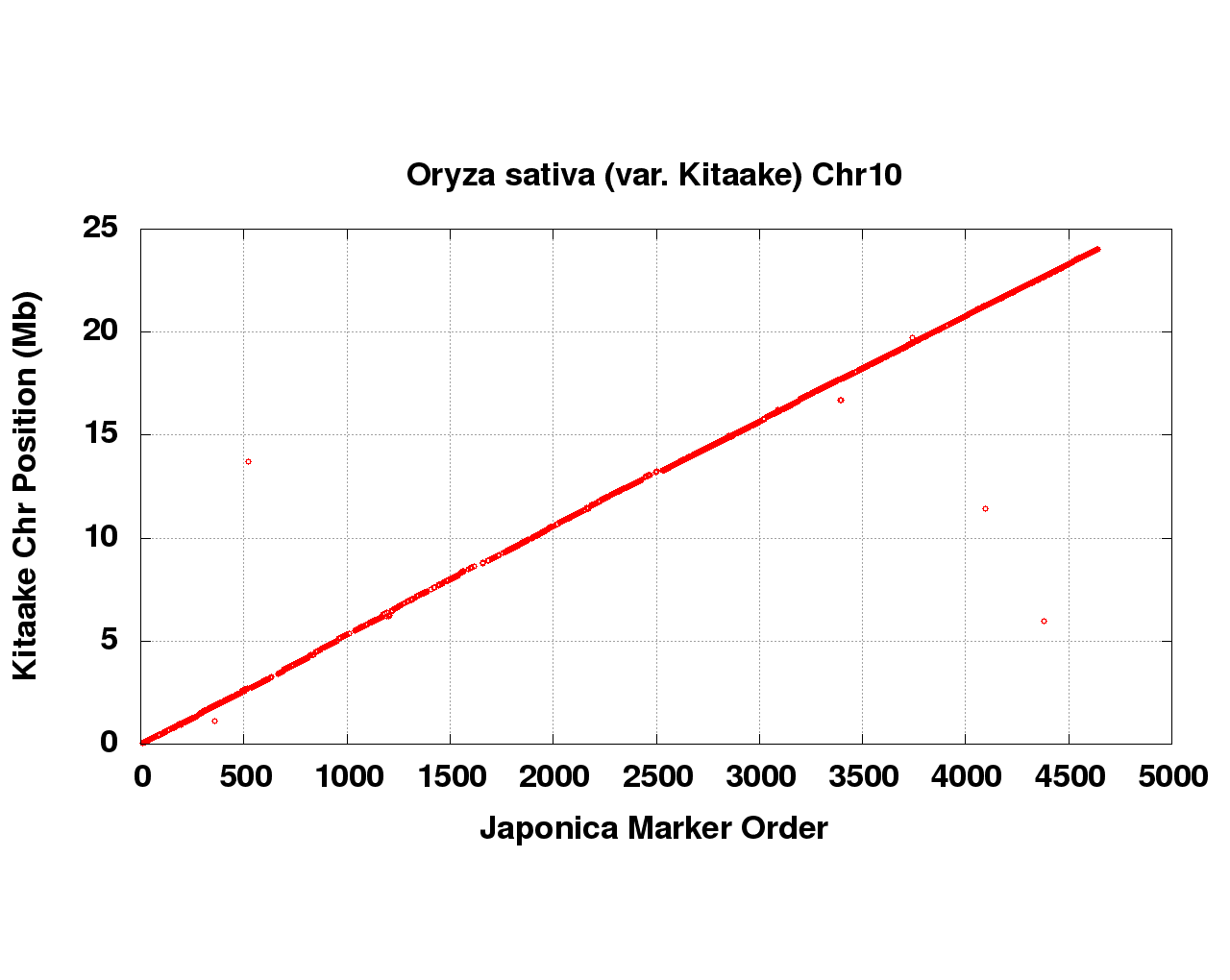

**Figure S10:** Syntenic Japonica sequence placements on the *Oryza sativa* (var. Kitaake) chromosome 10.

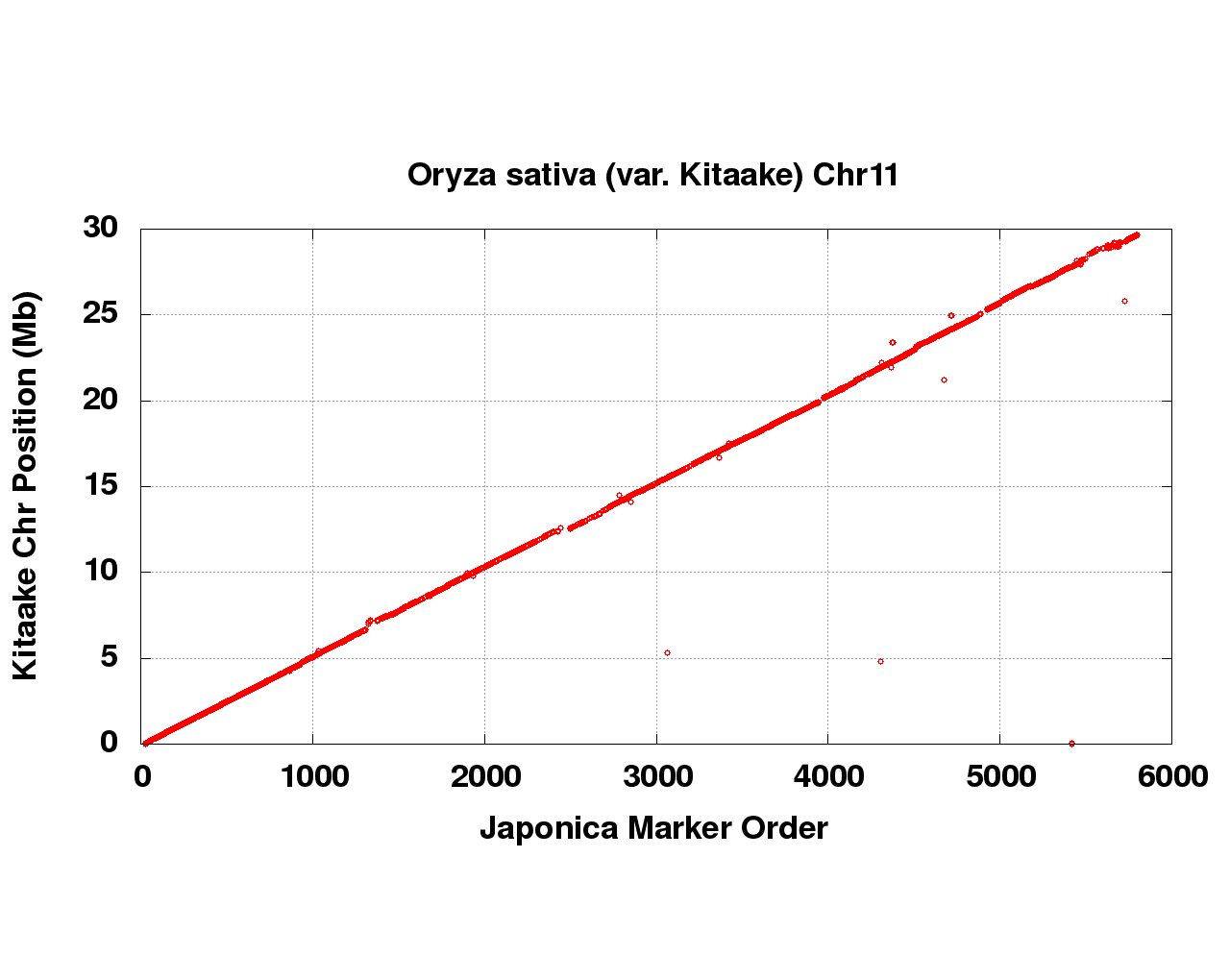

**Figure S11:** Syntenic Japonica sequence placements on the *Oryza sativa* (var. Kitaake) chromosome 11.

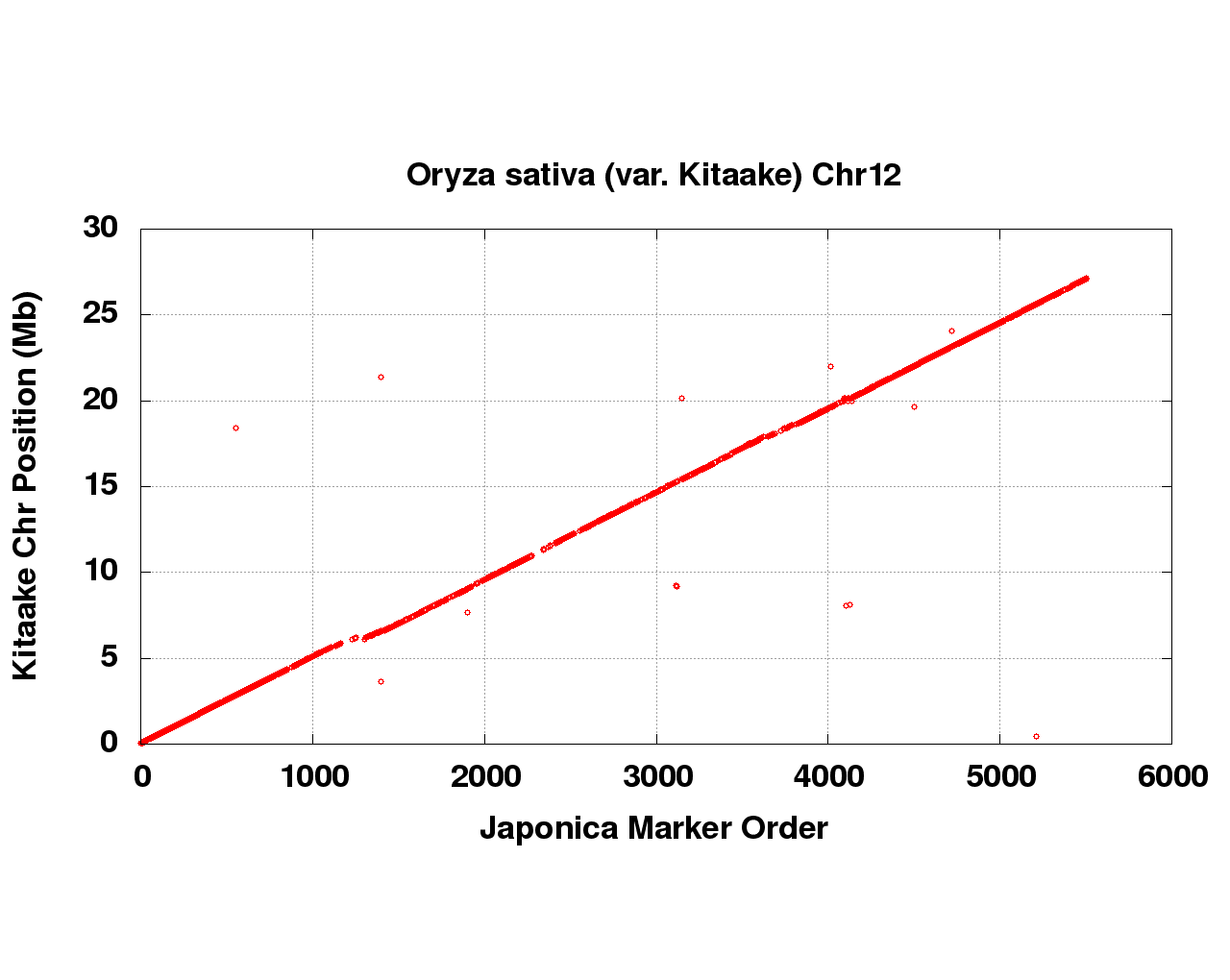

**Figure S12:** Syntenic Japonica sequence placements on the *Oryza sativa* (var. Kitaake) chromosome 12.

**S1.3 Screening and Final Assembly Release**

Scaffolds that were not anchored in a chromosome were classified into bins depending on sequence content. Contamination was identified using blastn against the NCBI nucleotide collection (NR/NT) and blastx using a set of known microbial proteins. Additional scaffolds were classified as repetitive (>95% masked with 24mers that occur more than 4 times in the genome) (67 scaffolds, 2.1 Mb), mitochondria (18 scaffolds, 800.2 Kb), prokaryotic (1 scaffolds, 20.9 Kb), chloroplast (7 scaffolds, 162.2 Kb), and low quality scaffolds with >50% of bases unpolished (8 scaffolds, 130.8 Kb). We also removed 15 scaffolds (529.5 Kb) representing the alternative haplotype of a section of sequence anchored in the chromosomes. Resulting final statistics are shown in Table S5.

| **Scaffold total** | 33 |
| --- | --- |
| **Contig total** | 476 |
| **Scaffold length total** | 381.6 Mb |
| **Chromosome Sequence** | 376.4 Mb |
| **Contig sequence total** | 377.6 Mb (1.0% gap) |
| **Scaffold N/L50** | 6 / 30.3 Mb |
| **Contig N/L50** | 75 / 1.4 Mb |

**Table S5**. Final summary assembly statistics for chromosome scale assembly.

**S1.3 Assessment of Assembly Accuracy:**

A set of 346 BAC clones was sequenced in order to assess the accuracy of the assembly. A range of variants was detected in the comparison of the BAC clones and the assembly. In 271 BAC clones, the alignments were of high quality (< 0.1% bp error); an example is given in Figure S13 (all dot plots were generated using Gepard [7]. Another 60 BACs indicate a higher error rate due mainly to their placement in repetitive regions (Figure. S14). The final 15 BAC clones are clones that indicate a rearrangement (10 clones) or a putative overlap on adjacent contigs (5 clones). Figure S15 is an example of a putative overlap on adjacent contigs within a chromosome. Typically, this result indicates that two local haplotypes have been assembled on the ends of adjacent contigs. It is common practice to collapse the two haplotypes into one haplotype unless they occur in repetitive areas where a collapse is more difficult to support. For this version of the genome, 22 adjacent contig pairs were detected that have overlapping ends (5 with clone support), but they were not collapsed because they occur in repetitive regions. The overall bp error rate (including marked gap bases) in the BAC clones is 0.09% (45,240 discrepant bp out of 46,414,368).

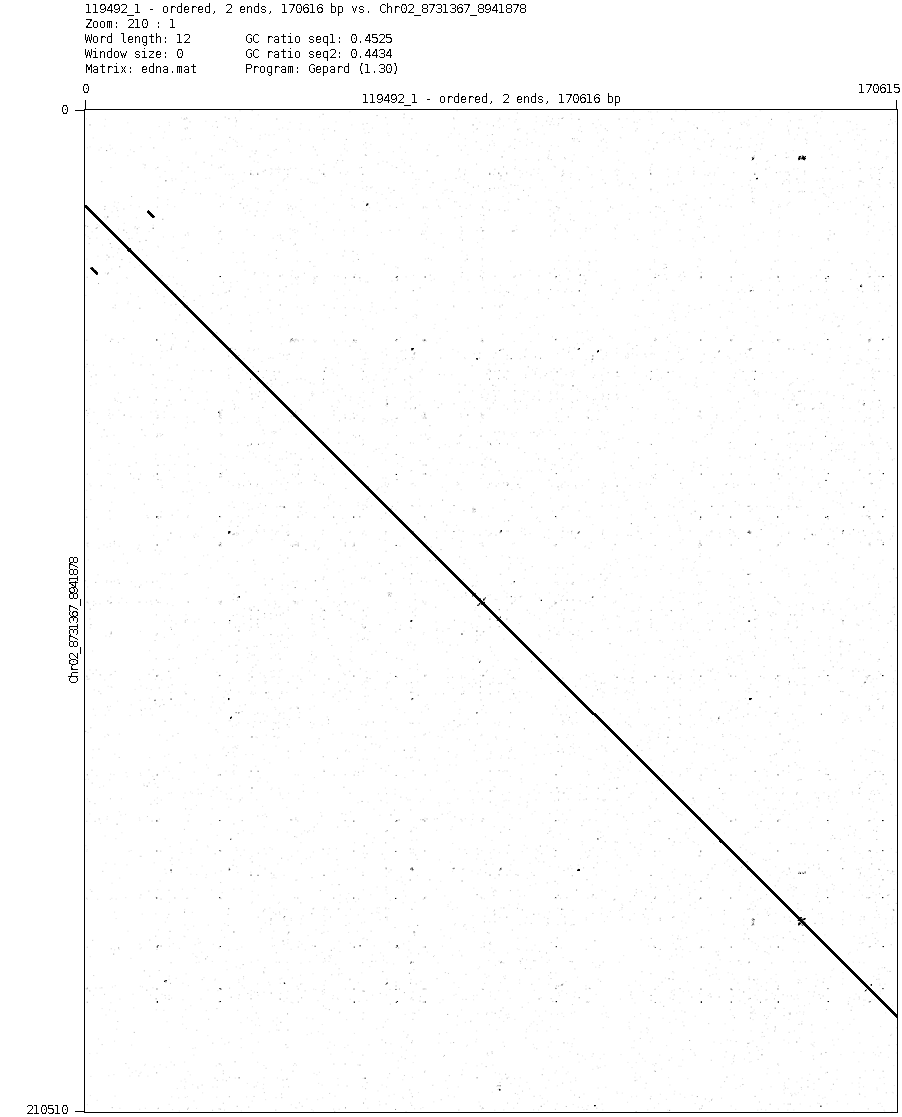

**Figure S13:** Dot plot of BAC clone 119492 on a region of Chr_02. This alignment is representative of the high quality BAC clone alignments in 271 of the 349 available BAC clones.

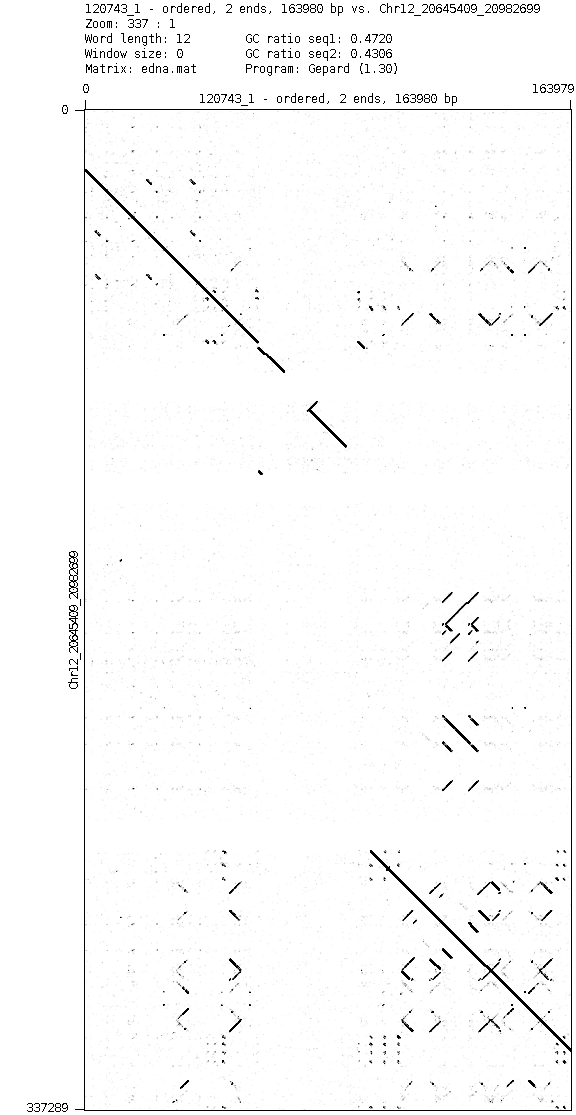

**Figure S14:** Dot plot of BAC clone 120743 on a region of Chr_12, which is representative of the 60 clones that landed in repetitive regions of the genome.

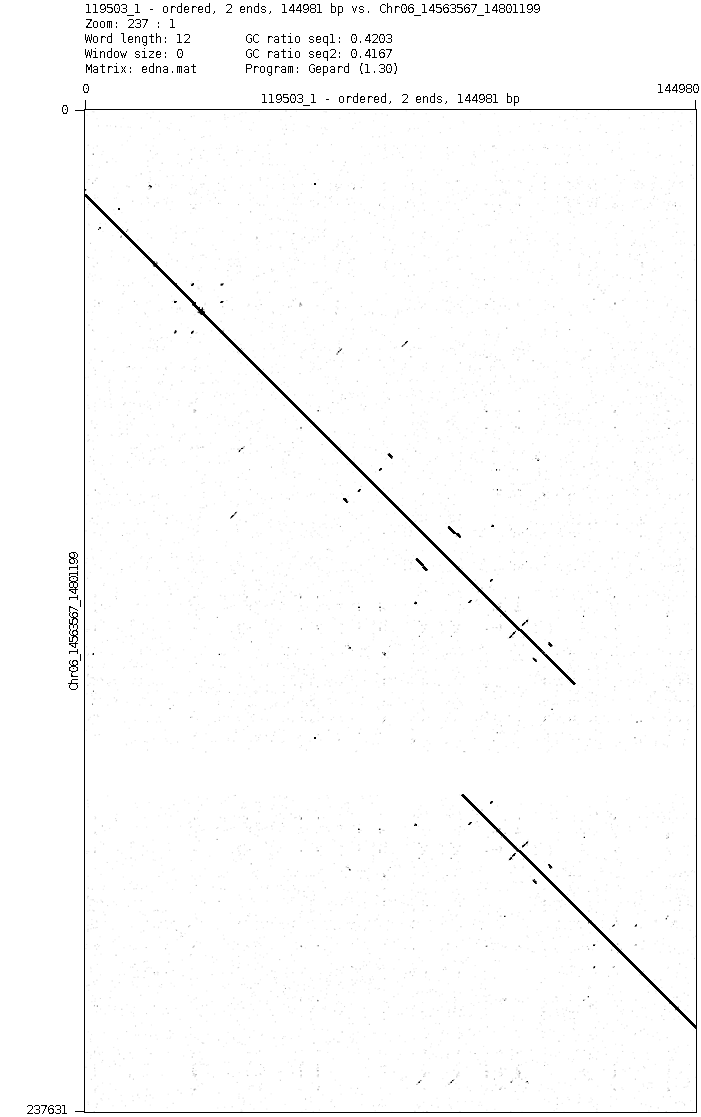

**Figure S15:** Dot plot of BAC clone 119503 in a region of Chr_06, representative of the 5 clones that align to an area where the adjacent contigs indicate that they overlap.

**BAC library construction:**

Arrayed BAC libraries were constructed using established protocols [8]. Briefly, young rice leaves (KitaakeX plants) were ground with liquid nitrogen and extracted for the isolation of intact nuclei. HMW DNA was isolated from nuclei imbedded in LMT agarose and lysed with detergent and proteinase K, followed by partial digestion with *Hind*III. After PFGE separation, appropriately sized HMW DNA fragments were cloned into the pAGIBAC1 vector [9]. Following mass transformation into DH10B T1 phage resistant *E coli* competent cells and spreading onto agar trays, BAC colonies were robotically picked into 384-well microtiter plates containing standard *E coli* growth media supplemented for long-term storage [8]. Similarly, a second Kitaake BAC library was constructed using the restriction enzyme *BstY*I. Copies of each BAC library were produced robotically and stored at -80^0^C.

| Lib name | Restriction enzyme | # colonies | Average insert size (kb) |
| --- | --- | --- | --- |
| OSJKBa | *Hind*III | 28,800 | 140 |
| OSJKBb | *BstY*I | 15,744 | 125 |

**Table S6:** Rice, KitaakeX, BAC libraries
