## Additional file 7 for "Genome sequence of the model rice variety KitaakeX"

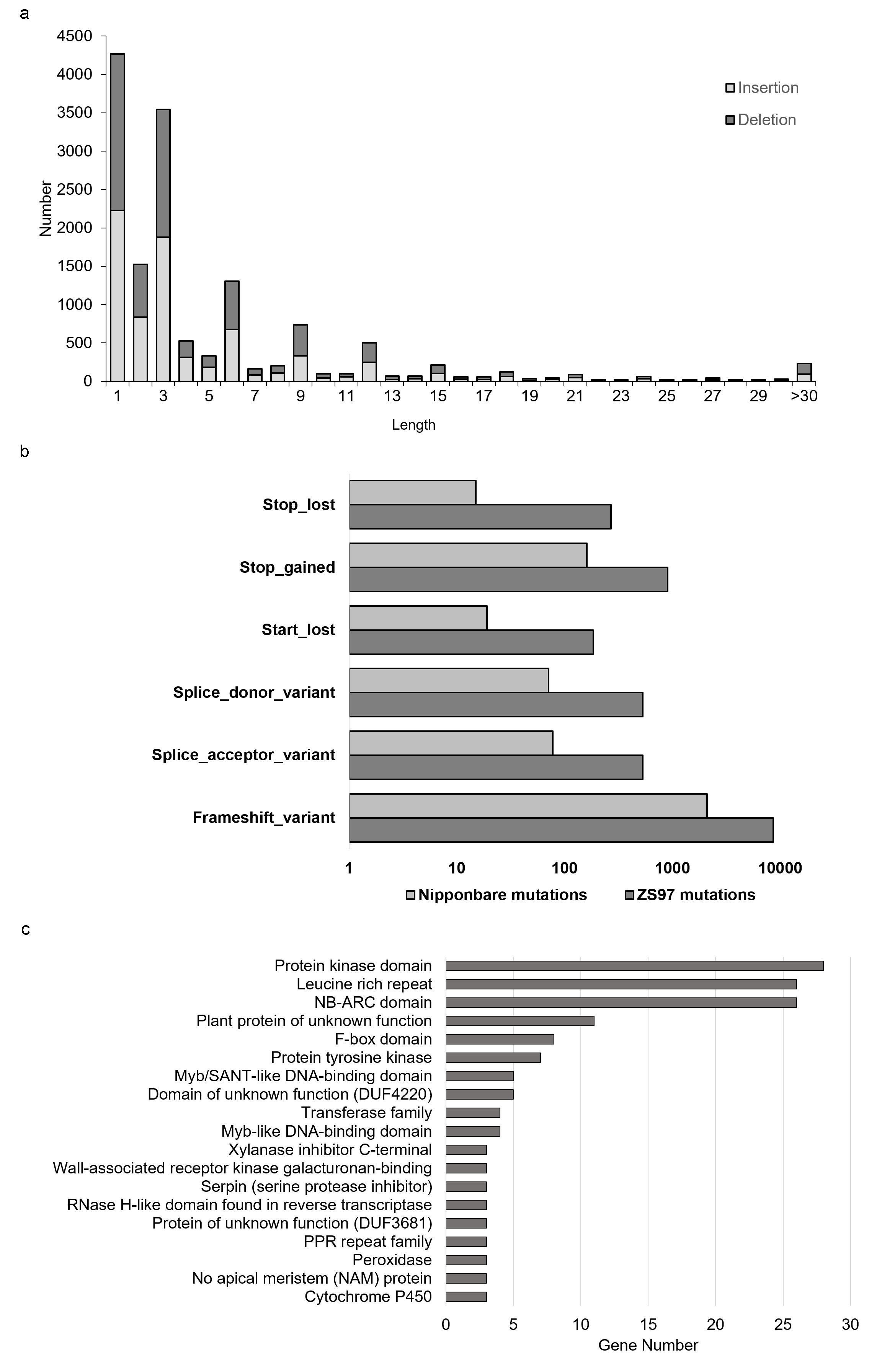
**Figure S16:** Genomic variation showing gene variation between KitaakeX and Nipponbare and ZS97. ***a*** Length distribution of InDels in protein-coding regions. The distribution indicates that nonsingle base InDels with a length of 3 bp (and/or multiples of 3) are much more abundant than the others. ***b*** SNPs and InDels that cause high-impact gene variations between KitaakeX and Nipponbare and ZS97. ***c*** Gene enrichment in KitaakeX unique present regions compared with Nipponbare.

pentatricopeptide repeat (PPR), NB-ARC (Nucleotide Binding)- (APAF-1 (apoptotic protease-activating factor-1), R proteins and CED-4 (*Caenorhabditis elegans* death-4 protein)),SANT domain (Swi3, Ada2, N-Cor, and TFIIIB).
