## Additional file 2 for "Genome sequence of the model rice variety KitaakeX"

|  | X.Kitaake | Nipponbare* | Zhenshan97* |
| --- | --- | --- | --- |
| Protein-coding genes | 35,594 | 39,045 | 34,610 |
| Complete BUSCOs (%) | 99.0 | 97.0 | 91.8 |
| Fragmented BUSCOs (%) | 0.3 | 1.0 | 0.8 |
| Missing BUSCOs (%) | 0.4 | 1.4 | 7.4 |

*Data are from Zhang et al (Zhang et al., 2017).

**Table S7**: Comparison of the X.Kitaake genome with other rice genomes.

| TE classification | X.Kitaake | | | | Nipponbare* | | | | Zhenshan97* | | | | |
| --- | --- | --- | --- | --- | --- | --- | --- | --- | --- | --- | --- | --- | --- |
|  | Length (kb) | | Percentage (%) | | Length (kb) | | Percentage (%) | | | Length (kb) | | Percentage (%) | |
| **Retrotransposons** | | 89,620 | | 23.49 | | 84,546 | | 22.59 | | | 92,184 | | 25.61 |
| Copia | 13,147 | | 3.45 | | 11,346 | | 3.03 | | | 9,462 | | 2.73 | |
| Gypsy | 70,797 | | 18.55 | | 66,689 | | 17.82 | | | 72,938 | | 21.03 | |
| LINEs | 3,992 | | 1.05 | | 3,350 | | 0.90 | | | 3,102 | | 0.86 | |
| SINEs | 374 | | 0.10 | | 1,426 | | 0.38 | | | 1,317 | | 0.37 | |
| Others | 1,310 | | 0.34 | | 1,735 | | 0.46 | | | 1,340 | | 0.37 | |
| **DNA transposons** | 32,552 | | 8.53 | | 63,146 | | 16.87 | | | 57,027 | | 15.84 | |
| Tc1-Mariner | 1,463 | | 0.38 | | 9,689 | | 2.59 | | | 8,940 | | 2.49 | |
| hAT | 1,792 | | 0.47 | | 4,337 | | 1.16 | | | 3,958 | | 1.10 | |
| Mutator | 7,484 | | 1.96 | | 15,736 | | 4.20 | | | 14,675 | | 4.08 | |
| PIF–Harbinger | 2,855 | | 0.75 | | 11,959 | | 3.19 | | | 11,126 | | 3.09 | |
| CACTA | 12,167 | | 3.19 | | 14,404 | | 3.85 | | | 11,925 | | 3.31 | |
| Helitron | 4,514 | | 1.18 | | 3,375 | | 0.90 | | | 3,081 | | 0.86 | |
| Others | 2,278 | | 0.59 | | 4,091 | | 1.09 | | | 11,695 | | 1.05 | |
| **Total** | 122,172 | | 32.02 | | 148,140 | | 39.58 | | | 149,662 | | 41.58 | |

*Data are from Zhang et al (Zhang et al., 2017).

**Table S8:** Summary of transposable elements in X.Kitaake, Nipponbare and Zhenshan97

|  | X.Kitaake vs. Nipponbare | X.Kitaake vs. ZS97 | Nipponbare vs.  ZS97* |
| --- | --- | --- | --- |
| SNPs | 253,295 | 2,328,319 | 2,665,280 |
| SNPs/kb | 0.67 | 6.17 | 7.14 |
| INDELs | 75,183 | 442,962 | 486,015 |
| INDELs/kb | 0.20 | 1.17 | 1.31 |

*Data are from the ZS97 genome paper (Zhang et al., 2016).

**Table S9:** Comparison of SNPs and INDELs between three rice genomes

| Substitution | X.Kitaake vs. Nipponbare | X.Kitaake vs. ZS97 | Nipponbare vs.  ZS97* |
| --- | --- | --- | --- |
| A->G | 43,744 | 390,891 | 447,240 |
| G->A | 42,497 | 436,480 | 497,244 |
| C->T | 42,375 | 434,854 | 495,844 |
| T->C | 44,114 | 392,605 | 449,693 |
| A->C | 10,004 | 87,253 | 99,660 |
| C->A | 10,566 | 91,047 | 103,398 |
| A->T | 11,150 | 93,603 | 110,671 |
| T->A | 11,597 | 95,297 | 112,538 |
| C->G | 8,049 | 63,856 | 72,939 |
| G->C | 8,538 | 64,714 | 73,645 |
| G->T | 10,092 | 90,079 | 102,695 |
| T->G | 10,569 | 87,640 | 99,713 |
| Transition | 172,730 | 1,654,830 | 1,890,021 |
| Transversion | 80,565 | 673,489 | 775,259 |
| Ti/Tv | 2.14 | 2.46 | 2.44 |

*Data are from the ZS97 genome paper (Zhang et al., 2016).

**Table S10:** Comparison of single base substitutions between three rice genomes

| Primary transcripts (loci) | 35,594 |
| --- | --- |
| Alternative transcripts | 12,900 |
| Total transcripts | 48,494 |
| **For primary transcripts:** |  |
| Average number of exons | 4.7 |
| Median exon length | 178 |
| Median intron length | 153 |
| **Gene model support (value is number of gene models):** |  |
| Any EST support | 31,854 |
| EST support over 100% of their lengths | 29,039 |
| EST support over 95% of their lengths | 29,511 |
| EST support over 90% of their lengths | 29,819 |
| EST support over 75% of their lengths | 30,334 |
| EST support over 50% of their lengths | 30,861 |
| Pfam annotation | 23,583 |
| Panther annotation | 23,142 |
| KOG annotation | 12,696 |
| KEGG Orthology annotation | 8,290 |
| E.C. number annotation | 8,948 |

**Table S11:** Oryza sativa Kitaake annotation v3.1 on assembly v3.0
