## Additional file 9 for "Genome sequence of the model rice variety KitaakeX"

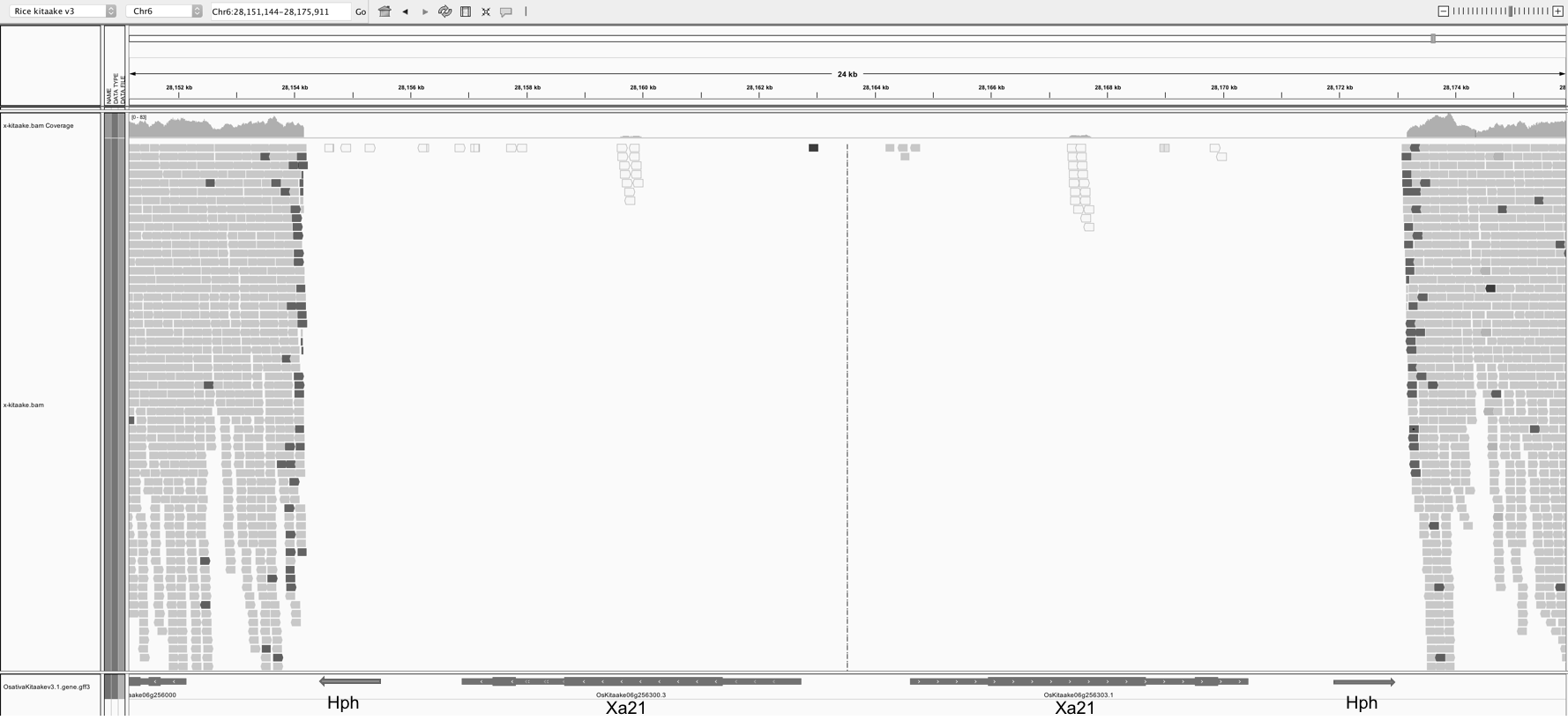

**Figure S17:** Integrative genomics viewer (IGV) snapshot showing presence of XA21 transgene and selectable marker encoding a hygromycin B phosphotransferase on chromosome 6 of KitaakeX.
